# Region-specific human brain organoids reveal synaptic and cell state drivers of glioblastoma invasion

**DOI:** 10.1101/2025.10.15.682580

**Authors:** Tarun N. Bhatia, Saisrinidhi Ganta, Mostafa Meselhe, Caitlin Sojka, Antoni Martija, Lisa Nieland, Uriel Rufen-Blanchette, Anson Sing, Alexia King, Brain Organoid Hub, Aparna Bhaduri, Kimberly Hoang, Edjah Nduom, Renee D. Read, Jeffrey Olson, Steven A. Sloan

## Abstract

Glioblastoma (GBM) is a highly heterogenous and malignant brain tumor, in part because it disrupts normal brain circuits to fuel its own growth and invasion. Thus, there is a need to identify the molecular features of invasive GBM cells and their regulators in the neural microenvironment. To address this in a fully human model, we engrafted patient-derived GBM cells (total *n=*15 independent samples) from three sources— fresh neurosurgical resections, cell lines, and whole GBM organoids—into human induced pluripotent stem cell-derived organoids patterned to forebrain, midbrain, and spinal cord identities. GBM cells from all sources infiltrated brain organoids within 2 days post-engraftment, reaching maximal invasion by day 14. Across organoids of distinct spatial and maturational milieu, GBM cells showed a consistent reduction in astrocyte-like states and an enrichment in neuron/glia progenitor-like (NPC-like) states. These NPC-like GBM cells expressed neuronal and synaptic machinery, and tumors enriched in this transcriptomic state prior to engraftment achieved greater organoid coverage, suggesting enhanced infiltration and synaptic integration of this GBM cell type. Although GBM cell states converged across organoid types after engraftment, infiltration was greater in the forebrain than spinal cord. This is likely reflective of synaptic input from deep-layer TBR1⁺ excitatory neurons in the forebrain, as demonstrated by a combination of rabies-based monosynaptic tracing and single-cell transcriptomics. In contrast, inhibitory neurons were the predominant synaptic partners of GBM in the spinal cord. Together, this fully human model of the neural-GBM connectome reveals how neuron-like GBM states and regionally distinct synaptic inputs cooperatively shape tumor invasion.

## INTRODUCTION

Histopathological studies from the 1930s revealed that glioma cells cluster around neuronal cell bodies^1,2^. These observations are consistent with recent work showing that glioma cells disrupt neural circuits, induce hyperexcitability in neurons, proliferate in response to neuron activity, and co-opt neuron-like mechanisms to invade brain tissue^3-9^. This bidirectional crosstalk between glioma cells and the nervous system is foundational to the emerging field of cancer neuroscience^10-12^. Seminal work in this field relied on rodent models and xenografts, but significant evolutionary differences at the level of cell types, topographic organization, and gene regulatory programs^13-15^ limit the ability of these models to fully recapitulate the human neural microenvironment. These differences are particularly pronounced in the cerebral cortex, the site of ∼90% of all adult gliomas^16^, which is larger and more complex in humans than in rodents due to an expanded pool of neural progenitors, such as radial glia and glial-intermediate progenitor cells^17,18^. These cells may either persist as quiescent remnants that are reactivated in adult tumors such as glioblastoma (GBM) or may reflect neurodevelopmental lineage programs that are hijacked by this highly aggressive tumor^19,20^. In addition to such progenitors, single-cell RNA sequencing has identified a spectrum of GBM cells that also reflect human neurodevelopmental cell types, including oligodendrocyte precursor (OPC)-like, astrocyte (AC)-like, mesenchymal (MES)-like, ciliated, and neuron-like cells^21-23^. These findings, along with the capacity of GBM to transition between transcriptomic states in response to microenvironmental cues, emphasizes the need to study GBM behaviors within a fully human neural niche.

Thus, groups are increasingly adopting three-dimensional (3D) organoid-based models, which facilitate the use of GBM cells directly from neurosurgical tissues^24^. These 3D models, including whole GBM organoids^25-27^, tumor explants^28^, and matrix-embedded cultures,^29-31^ preserve GBM-intrinsic features from the patient tumor, but lack the host neural microenvironment. To address this limitation, human brain organoids derived from embryonic stem cells (hESCs) or induced pluripotent stem cells (hiPSCs) have been used. Key advantages of these 3D brain organoids are that their microenvironment can be precisely manipulated, and they recapitulate aspects of the human brain cytoarchitecture, permitting interrogation of GBM migration and integration into neural circuits. While early approaches used genetically engineered brain organoids to model gliomagenesis^32,33^, recent efforts involve engrafting cells from gliomasphere cell lines or fresh surgical resections into wildtype hESC/hiPSC-derived brain organoids^20,34-43^. These fully human-based models, referred to as GLICO^35,36,38,41^, HOTT^39,40,42^, or GCO^43^ systems maintain the genetic and cell type diversity of the patient tumor within a human neural context. Using unpatterned cerebral organoids or dorsal forebrain-patterned cortical organoids, these models^20,34-43^ have provided insights into the intrinsic and extrinsic mechanisms underlying GBM cell diversity, infiltration, intercellular transfer, GBM cell plasticity, and GBM cell responses to therapeutic agents.

Another unique advantage to brain organoids is that they can be regionally patterned to mimic distinct areas of the human CNS^44-51^ and matured for months in culture, mirroring the *in vivo* developmental trajectory of neural cells^49-51^. These features provide a unique opportunity to unmask how host regional identity and cell type composition influence the proliferation, infiltration, and transcriptomic signatures of engrafted GBM cells. This question is relevant given recent findings that glioma cells express receptors for many neurotransmitters and receive synaptic input from diverse neuron populations, including long-range projections from local and extracortical regions^4,52-54^. We therefore investigated if GBM cells adopt unique or universal transcriptomic signatures after entry into the distinct microenvironments offered by forebrain, midbrain, and spinal cord organoids. To accomplish this, we engrafted tumor cells from the same GBM resection into these distinct neural niches under tightly matched experimental conditions. This design, enabled by the Brain Organoid Hub’s routine production of ready-to-engraft region-specific brain organoids generated simultaneously from the same hiPSC lines and differentiations minimizes variability and allows direct comparison of tumor behaviors across distinct human neural contexts.

Beyond tumor-intrinsic transcriptional programs, GBM infiltration may also depend upon the ability of GBM cells to engage functionally with region-specific host neurons through synaptic input and paracrine cues. This possibility is clinically significant because although GBM initiation is more common in the forebrain, infiltration into the hindbrain is a frequent observation at end-stages^55^. This pattern suggests that region-dependent cues may promote or restrict tumor invasion. Thus, we also investigated spatial biases in GBM infiltration and mapped presynaptic partners of GBM after entry into regionally patterned organoids, using a combination of rabies-based monosynaptic tracing and single-cell transcriptomics. Together, our work uses a fully human platform to dissect the neuron-tumor connectome across spatial and maturational milieu, enabling the identification of circuit-level interactions that drive GBM synapse formation, shape tumor cell state signatures, and ultimately promote infiltration.

## RESULTS

### Glioblastoma (GBM) cells infiltrate brain organoids within 2 days post-engraftment

To understand how GBM infiltrates neural tissues and integrates into neural circuitry, we needed a system that incorporates patient-derived tumors into a human neural context. We accomplished this by using a modified version of the GLICO^35,36,38,41^, HOTT^39,40,42^, and GCO^43^ systems. We dissociated neurosurgically resected GBM (**Supplementary Table 1**), labeled cells with a GFP-expressing lentivirus, and engrafted them into wildtype human iPSC-derived region-specific brain organoids (**Fig. 1A**). This presented several optimization challenges. First, we needed to minimize the time from operating room to organoid. Second, we wanted to preserve as much cell type heterogeneity of the parent tumor as possible. Third, we needed to maximize the number of GBM cells labeled with GFP prior to engraftment. We tested 3 separate approaches to culture and label dissociated tumor cells in this *in vitro* setting (**Fig. 1A**): (1) tumor cells maintained in suspension in an Eppendorf tube and labeled with the GFP-lentivirus for 4 hours on a shaker, (2) tumor cells in suspension in a tube and labeled with the GFP-lentivirus overnight (∼18 hours) on a shaker, and (3) tumor cells plated in 2D on Matrigel and exposed to the GFP-lentivirus for 24 hours.

**Fig. 1.**
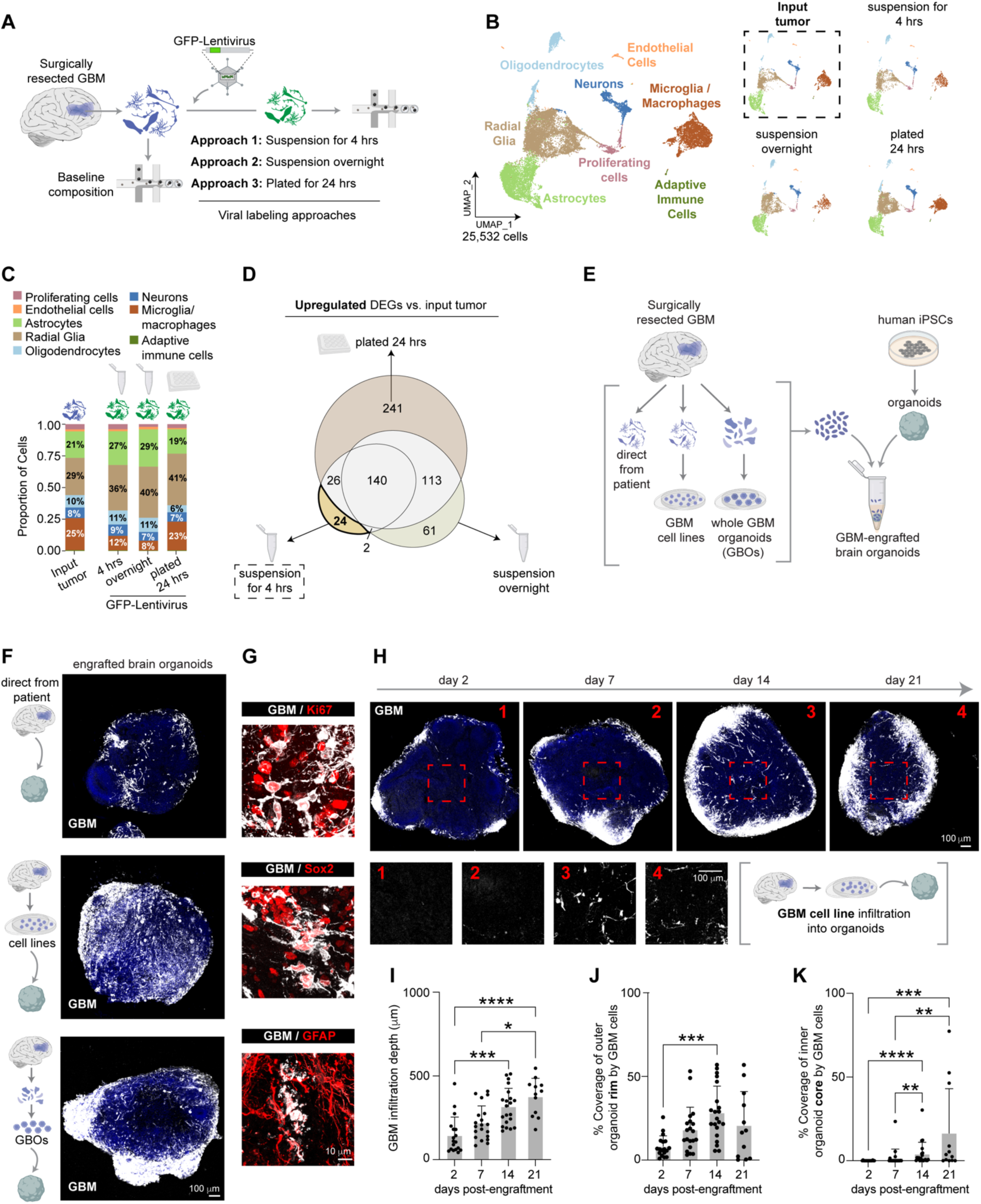
Glioblastoma (GBM) cells infiltrate brain organoids within 2 days post-engraftment. (**A**) Surgically resected GBM tissues were dissociated into a single-cell suspension and labelled with a GFP-lentivirus (pLenti-CMV-GFP) at a multiplicity of infection (MOI) of 5 by exposing cells to the virus for 4 hours in suspension culture on a shaker (Approach 1), overnight in suspension culture on a shaker (Approach 2), or by plating cells in 2D on Matrigel and exposing cells to the virus for 24 hours (Approach 3). Single-cell RNA sequencing was performed following lentiviral exposure and compared to single-cell transcriptomes following dissociation of the input tumor. (**B**) UMAP of 25,532 individual cells that passed QC and assigned to clusters, annotated based on expression of cell type-specific markers. (**C**) Proportion of cells in each of the four samples. (**D**) Area-proportional Venn diagram to visualize the overlap between differentially upregulated genes when each of the lentiviral-exposed samples were compared to the dissociated, input tumor. The short, 4-hour GFP-lentivirus exposed GBM cells maintained in suspension culture showed the fewest differentially upregulated genes (24). (**E**) Schematic illustration of engraftment of GBM cells from 3 sources—harvested directly from neurosurgical resections, patient-derived GBM cell lines, and whole GBM organoids (GBOs)—into human iPSC (hiPSC)-derived brain organoids. (**F**) Representative images of GFP^+^ GBM cells (white) from neurosurgical resections (direct-from-patient), patient-derived cell lines, or whole GBOs engrafted into hiPSC-derived brain organoids. (**G**) High-magnification images showing direct-from-patient GBM cells (white) expressing canonical markers of stemness (Ki67 and Sox2; red) and glia (GFAP; red) within organoids. (**H**) Representative low-magnification (top row) and high-magnification (bottom row) images of GBM cells from patient-derived GBM cell lines across days 2 to 21 post-engraftment into brain organoids. (**I**) Quantification of the infiltration depth of patient-derived cell lines after entry into brain organoid sections, across days 2 to 21 post-engraftment. For **I**, each dot represents the average infiltration depth of 5 randomly selected GBM cells in the inner core of the brain organoid section, calculated as the shortest distance from each cell to the edge of the organoid. (**J**) Quantification of the area of the outer 25% (“rim”) of brain organoid sections covered by patient-derived cell lines, across days 2 to 21 post-engraftment. (**K**) Quantification of the area of the inner 25% (“core”) of brain organoid sections covered by patient-derived cell lines, across days 2 to 21 post-engraftment. For **J-K**, each dot is the area coverage in a single organoid section. Data in **I-K** are from two independent GBM cell lines engrafted into two independent hiPSC-derived brain organoid lines (*n=*2 GBM lines x 2 organoid lines). Data in **I-K** were non-normal (per Shapiro-Wilk) and analyzed by Kruskal-Wallis followed by the Dunn’s test to control for multiple comparisons. All error bars are mean + SD.

To test which culture and labeling approach best preserved host tumor biology, we performed single-cell RNA sequencing on tumor cells following each lentiviral exposure scheme and compared the results to those from the input, dissociated parent tumor prior to lentiviral exposure (**Fig. 1B**). We manually annotated clusters based on expression of neural and immune cell type-specific marker genes (**Fig. S1A-C**). Reassuringly, expected GBM cell types were broadly preserved across lentiviral exposure and culture conditions (**Fig. 1B-C**). However, differential gene expression (DEG) analyses revealed that GBM cells exposed to GFP-lentivirus for 4 hours in suspension showed the *fewest* uniquely upregulated genes (only 24; **Fig. 1D**; **Supplementary Table 2**) compared to the parent tumor, and these genes were related to the cytoskeleton (**Fig. S1D**). This group also showed the fewest uniquely downregulated genes (17; **Fig. S1E**; **Supplementary Table 2**), related to stress and immune response pathways (**Fig. S1F**). Longer exposures of GBM to the GFP-lentivirus showed significantly greater DEGs relative to the parent tumor (**Fig. 1D**; **Fig. S1D-F**), including alterations in pathways linked to neurogenesis and gliogenesis (**Fig. S1F**). Thus, despite observing robust infiltration of GFP^+^ GBM cells into organoids under all tested culture conditions (GBM cells pseudocolored white in **Fig. S1G**), we proceeded with a short, 4-hour exposure of suspended GBM cells to a GFP-lentivirus prior to engraftment, as this best maintained the cell composition and transcriptional profile of the input, parent tumor.

After attempting many physical engraftment strategies, we achieved the most reproducible success by immersing an organoid in a suspension of ∼50,000-100,000 GFP-labeled GBM cells in a v-bottom (conical) tube for ∼36-48 hours before transferring to an ultra-low attachment plate for the remainder of the experiment. Much of our work relied on using GBM cells directly harvested from patients. To account for the inherent unpredictability of obtaining viable tumors, we validated our findings with additional GBM models, including dissociated patient-derived gliomasphere cell lines and dissociated whole glioblastoma organoids (GBOs) generated from minced patient tissues (total *n=*15 independent patient samples; **Fig. 1E**; **Supplementary Table 1**). GFP^+^ GBM cells (pseudocolored white in **Fig. 1F-G**) from all three settings (directly from surgical resections, dissociated patient-derived gliomaspheres, dissociated GBOs) infiltrated brain organoids and maintained expression of stemness (Ki67, Sox2) and glial markers (GFAP).

How long does it take for GBM cells to infiltrate a 2-3mm thick organoid? To answer this question, we examined the temporal dynamics of GBM invasion by quantifying the infiltration depth of GBM cells and assessing their spatial coverage within the outer rim vs. inner core of brain organoids (**Fig. 1H-K**). GBM cells begin to enter brain organoids as early as 2 days post-engraftment but were restricted to the outer rim at this early time-point. After 14 days GBM cells had often infiltrated the inner core of organoids. Based on this finding, we performed our downstream transcriptomic analyses at 14 days post-engraftment.

### GBM cells from neurosurgically resected tissues are enriched in a neuron/glia progenitor-like (NPC-like) transcriptomic state post-engraftment into regionally patterned organoids

After validating the GBM-engraftment protocol in our hands, we assessed how the distinct spatial context provided by region-specific brain organoids alters the transcriptomic state of the tumor. We engrafted cells from the same GBM patient sample (*n=*3 samples; **Supplementary Table 1**) into dorsal forebrain, ventral forebrain, midbrain, and spinal cord organoids, all derived simultaneously from the same hiPSC lines and differentiations to reduce batch variability (**Fig. 2A**). These four brain regions span a wide neuroanatomical spectrum across the major vesicles of the CNS and are distinct in cell type composition and signaling cues, providing a diverse set of neural niches for matched comparisons. We used established protocols^44-48^ to pattern organoids to these regions and validated regionalization at transcriptomic (**Fig. S2**) and protein (**Fig. 2B**; top row) levels.

**Fig. 2.**
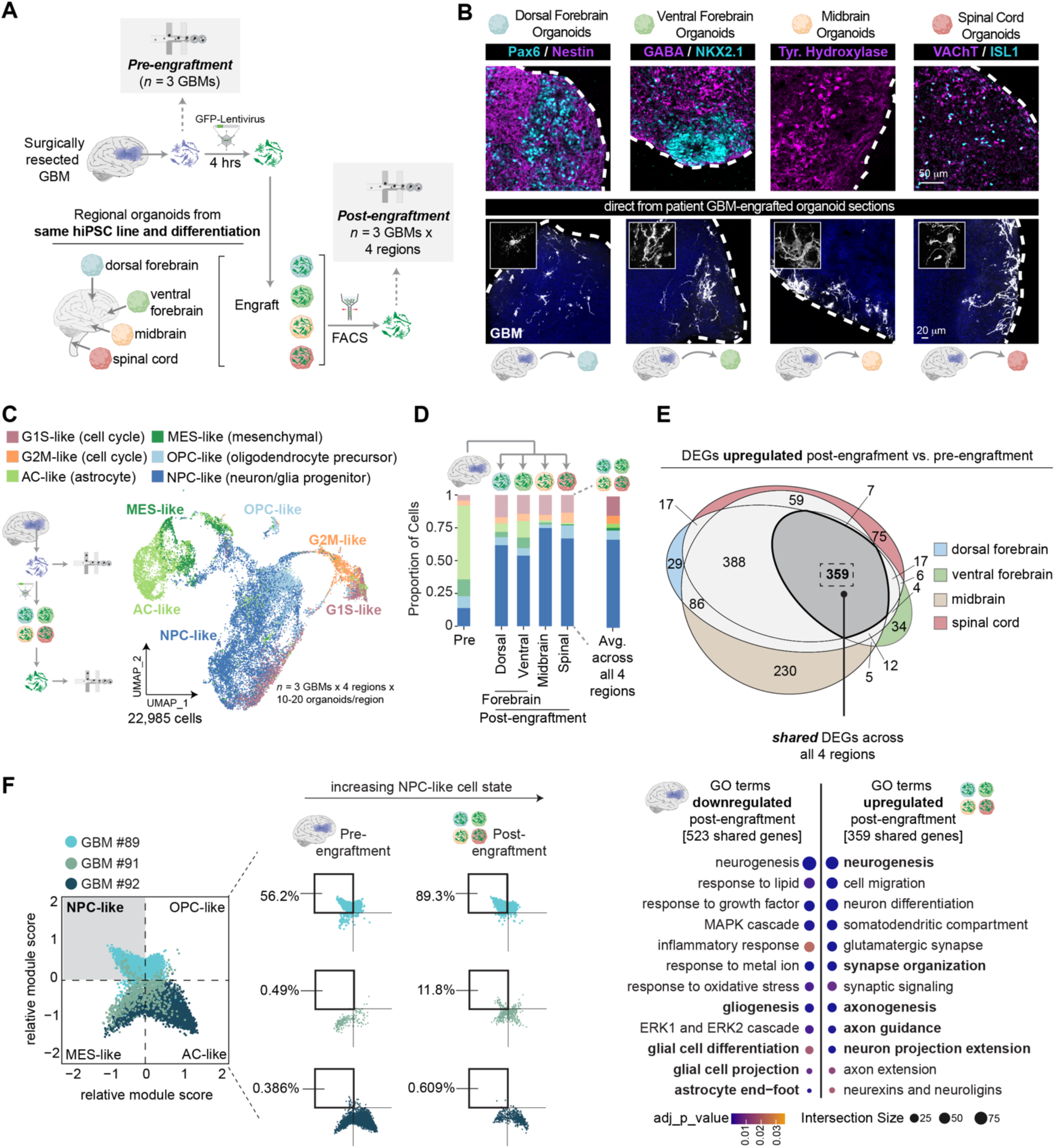
GBM cells converge onto a neuron/glia progenitor-like (NPC-like) transcriptomic state after engraftment into forebrain, midbrain, and spinal cord organoids. (**A**) GBM cells from surgical resections (*n=*3 independent patient samples) were dissociated, labeled with a GFP-lentivirus for 4 hours in suspension culture, and then engrafted into dorsal forebrain, ventral forebrain, midbrain, and spinal cord organoids all derived in parallel from the same hiPSC lines and differentiations to reduce batch variability. After 14 days of organoid engraftment, GFP^+^ GBM cells were isolated via FACS and captured for single-cell RNA sequencing. For every post-engraftment GBM there was a matched pre-engraftment GBM to track the changes in the transcriptomic signatures of GBM cells after entry into the organoid niche. (**B**; top row) Representative images of dorsal forebrain organoids stained for Pax6 (cyan) and Nestin (magenta), ventral forebrain organoids stained for NKX2.1 (cyan) and GABA (magenta), midbrain organoids stained for tyrosine hydroxylase (red), and spinal cord organoids stained for ISL1 (cyan) and VAChT (magenta). (**B**; bottom row) Representative images of direct-from-patient GBM cells (white) infiltrating dorsal forebrain, ventral forebrain, midbrain, and spinal cord organoids at low magnification and high magnification (insets; bottom row). (**C**) UMAP of 22,985 individual malignant cells that passed QC and after exclusion of non-malignant cells based on copy number variations. Malignant cells were assigned to clusters based on Neftel *et. al.*, 2019^21^. (**D**) Proportion of cells across the six Neftel GBM cell types in each of the four region-specific brain organoids and in the parent tumor. (**E**) Area-proportional Venn diagram to visualize the overlap between differentially upregulated genes when GBM cells engrafted into each regional organoid were compared to the dissociated tumor pre-engraftment. A majority of differentially upregulated genes (359) were shared across all four regional microenvironments. Select gene ontology (GO) terms from gProfiler reflecting the 359 genes that are upregulated in GBM cells engrafted into all four regional organoids, compared to the GO terms over-represented among the shared 523 genes that are downregulated post-engraftment into the four regional organoids. (**F**) Two-dimensional representation of GBM states pre-engraftment vs. post-engraftment across each of the three independent GBM samples.

GBM cells successfully infiltrated all four region-specific organoids and retained their morphological complexities within the neural microenvironment (**Fig. 2B**; bottom row). We applied FACS to isolate GFP^+^ cells (GBM cells) 14 days after engraftment and performed single-cell RNA sequencing on post-engrafted cells. These transcriptional profiles were compared to matched pre-engraftment GBM samples, to control for tumor-intrinsic differences across patients and reveal true microenvironment-driven changes. A total of 42,483 cells passed quality control (**Fig. S3A-B**) and cluster annotations revealed the presence of all expected neural cell populations (**Fig. S3B-C**). We profiled for copy number variations (CNVs) using our established pipeline^50^ to exclude non-malignant cells, which may have been included as a result of FACS artifacts, infiltrating cells from the tumor margin, or, as recently suggested^43^, as a result of transfer of GFP transcripts from malignant to non-malignant cells (**Fig. S3D-J**). In brief, this pipeline involved correlating CNV signals of each cell in our dataset with the average CNV signal across all cells from the same tissue. As a second method, we validated this pipeline by correlating expression of *EGFP* in our post-engraftment samples with CNVs, observing a direct link (*r=*0.5060; **Fig. S3G-H**). Based on this pipeline, we confirmed that most of the input tumor pre-engraftment was malignant (70.1%), a proportion maintained among EGFP^+^ cells post-engraftment (**Fig. S3I-J)**. As expected, the EGFP^-^ cells post-engraftment (likely false positives from FACS) were typically classified as non-malignant per our computational CNV analysis.

We then annotated the malignant clusters as G1S-like (cell cycle), G2M-like (cell cycle), astrocyte (AC)-like, mesenchymal (MES)-like, oligodendrocyte precursor (OPC)-like, and neuron/glia progenitor (NPC)-like, based on Neftel *et. al.*, 2019^21^. After engraftment into organoids, tumor cells converged onto an NPC-like state regardless of regional organoid identity (**Fig. 2C-D**; 67.2% NPC-like cells across all 4 regions post-engraftment versus 13.9% in the input tumor). This enrichment in the NPC-like state was accompanied by a *downregulation* in the astrocyte-like state, which was typically the dominant signature in the input tumor (**Fig. 2D**; 56.2% astrocyte-like cells in the input tumor versus 3.2% astrocyte-like cells post-engraftment across all 4 regions). When we performed DEG analyses between pre- and post-engraftment conditions in each region, we surprisingly found that most post-engraftment changes were shared across all four regions (**Fig. 2E**; **Fig. S4A-C**; **Supplementary Table 3**). GO analyses of the 359 genes commonly upregulated in GBM across all four regions revealed processes related to neurogenesis, migration, synaptic signaling, axon guidance, and developmental signals such as neurexins-neuroligins (**Fig. 2E**). In contrast, 523 genes were commonly downregulated in GBM across all four regions and were related to gliogenesis, glial projection, and inflammatory or stress response pathways (**Fig. 2E**; **Fig. S4C**; **Supplementary Table 3**). Thus, the NPC-like state is a shared transcriptomic state among GBM cells across brain regions and this observation was consistent across all three GBM tissues, albeit the effect size varied by sample (**Fig. 2F**). This underscores the contribution of intrinsic factors such as genetic background in addition to extrinsic cues.

### GBM gliomasphere cell lines are also enriched in NPC-like signatures post-engraftment

We next asked if the enrichment of the NPC-like state was also evident when dissociated gliomaspheres from patient-derived cell lines are engrafted into neural organoids. For these experiments, we engrafted tumor cells into organoids of the opposite sex, allowing us to use the expression of Y-chromosome-specific genes to further validate our CNV-based computational pipeline (**Fig. S5A-E**). Consistent with our previous observations, malignant gliomasphere cells also converged onto the NPC-like state within organoids representing all four brain regions (**Fig. S5F-G**). Given that GBM cells adapt to their microenvironment and rely on neighboring neural cells to propel their own growth^3-5,8,9,56^, we also asked whether this universal convergence on the NPC-like state is mediated by shared extrinsic cues across all four regions. We performed single-cell RNA sequencing on GFP-negative cells (the “host” organoid fraction) following FACS isolation of gliomasphere-engrafted organoids and used the cell-cell communication pipeline, CellChat^57^, to identify putative host-secreted ligands that may influence tumor cell states post-engraftment. This analysis revealed many shared ligands such as PTN, CNTN, NCAM, across all four regions (**Fig. S5H-K**; region-specific exceptions in bold) that belong to developmental signaling pathways and are consistent with prior work identifying tumor-extrinsic cues that influence GBM^42,58,59^.

We next asked if the transcriptomic changes observed when gliomasphere cells are engrafted into organoids are driven by the neural microenvironment or can be induced under simplified culture conditions lacking growth factors such as EGF and FGF2. To test this, we performed bulk RNA sequencing on gliomaspheres and on the same cells cultured in monolayer for 14 days in forebrain, midbrain, or spinal cord organoid media. Transcriptomically, GBM cells exhibited molecular signatures dominated primarily by cell line and secondarily by culture conditions (3D gliomaspheres or 2D adherent), irrespective of the media (**Fig. S6A-B**). We then compared the differentially upregulated genes between gliomasphere cells cultured in 2D in forebrain media to those after engraftment into dorsal forebrain organoids. Only a small fraction of genes (*n=*40; **Supplementary Table 4**) overlapped, suggesting largely unique phenotypic shifts (**Fig. S6C-D**; 40 intersecting genes highlighted in the volcano plot), driven by activation of developmental signaling pathways (Notch, NTRK) within the organoid niche. Notably, DEG analyses also revealed that most differentially expressed genes were shared across cells plated in different regional organoid media (**Fig. S6E-F**), consistent with lack of region-specific transcriptomic changes when GBM cells were engrafted into organoids (**Fig. 2** and **Fig. S5**)

### The NPC-like state persists in GBM cells engrafted into more mature dorsal forebrain organoids

We asked whether enrichment of the NPC-like state depends on host developmental stage by engrafting GBM cells into immature, neurogenic dorsal forebrain organoids (∼day 36-64 at engraftment) or more mature, gliogenic organoids (∼day 208-279 at engraftment), and comparing transcriptomic identities (**Fig. 3A**). We used dorsal forebrain organoids for these experiments as our group has previously characterized the temporal dynamics of neurogenesis and gliogenesis within this region^50,51^ (**Fig. 3B**). Surgically resected GBM cells infiltrated dorsal forebrain organoids across developmental stages (**Fig. 3C**). We annotated malignant clusters and again observed increases in NPC-like signatures within immature and mature microenvironments across all three GBM samples (**Fig. 3D-F**). Interestingly, the effect size was slightly diminished in more mature microenvironments (**Fig. 3E**; 30.53% NPC-like GBM cells in mature organoids versus 43.48% NPC-like GBM cells in immature organoids). DEG analyses revealed that while the majority of differentially upregulated genes post-engraftment were shared across both microenvironments (187 genes; **Supplementary Table 5**), a substantial proportion of genes were also unique (134 genes unique to immature and 141 genes unique to mature organoids; **Fig. 3G**; **Supplementary Table 5**). Genes common to both developmental ages were related to neurogenesis, migration, axon guidance, and notch signaling (**Fig. 3G**). However, genes related to cholesterol and steroid biosynthesis were specifically enriched in GBM engrafted into immature microenvironments, whereas genes related to vascular endothelial growth factor (VEGF) signaling and insulin-like growth factor (IGF) signaling were enriched in more mature milieu (**Fig. 3H** and **Fig. S7A-B**). Importantly, gliomasphere cells also showed an increase in NPC-like states after entry into mature dorsal forebrain organoids, independent of their genetic background (**Fig. S8**). These data suggest that although the same NPC-like state is enriched across developmental ages, the signaling pathways underlying this cell state enrichment differ in immature vs. mature environments, reflecting the impact of distinct host cell types present throughout organoid development.

**Fig. 3.**
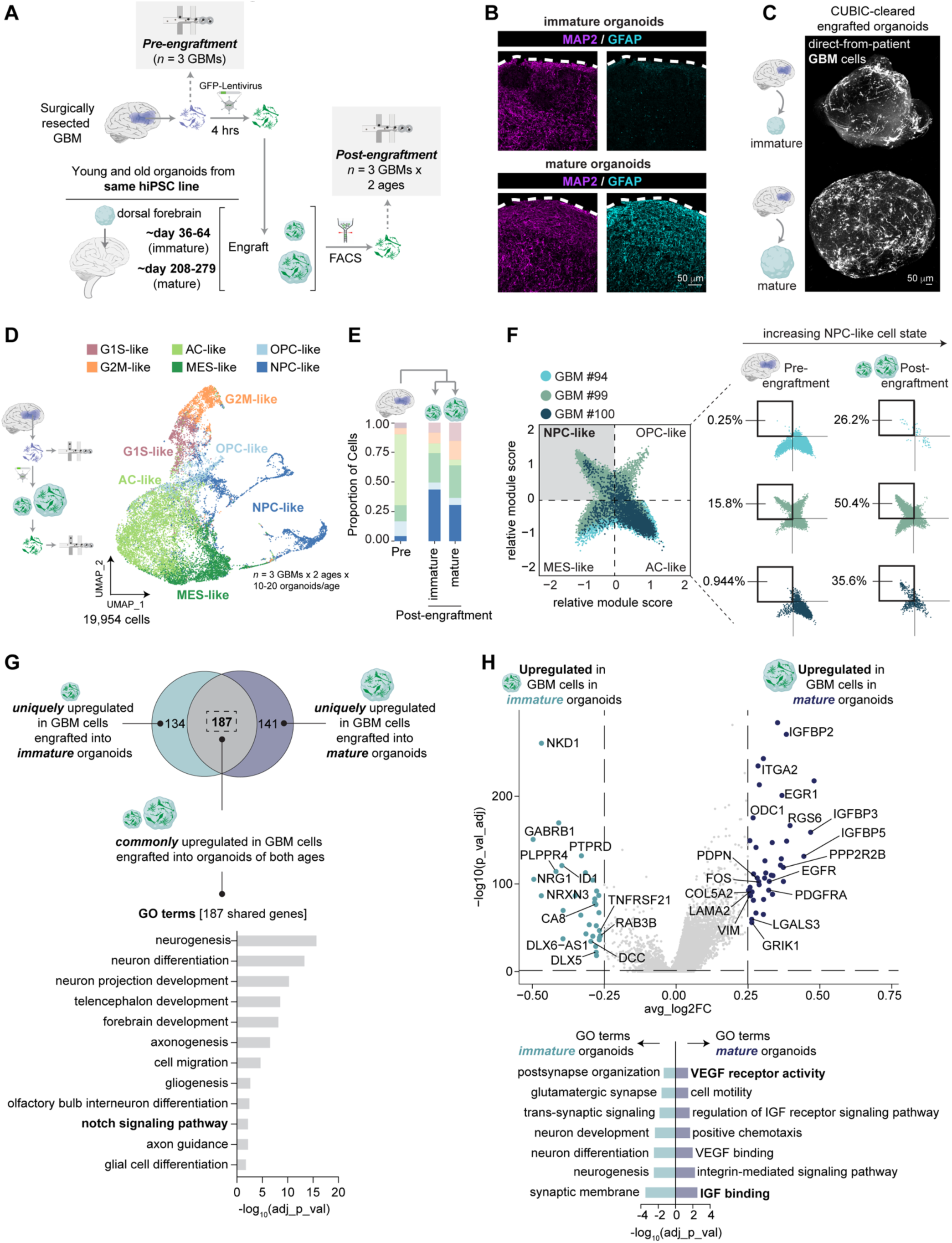
The NPC-like GBM transcriptomic state persists after engraftment to more mature (day 200+) dorsal forebrain organoids. (**A**) GBM cells from surgical resections (*n=*3 independent patient samples) were dissociated, labeled with a GFP-lentivirus for 4 hours in suspension culture, and then engrafted into dorsal forebrain organoids that were immature (∼day 36-64 on the day of engraftment) vs. dorsal forebrain organoids that were more mature (∼day 208-279). After 14 days of organoid engraftment, GFP^+^ GBM cells were isolated via FACS and captured for single-cell RNA sequencing. For every post-engraftment GBM there was a matched pre-engraftment GBM to track the changes in the transcriptomic signatures of GBM cells after entry into the organoid niche. (**B**) Representative images of immature vs. mature dorsal forebrain organoids stained for neuron marker MAP2 (magenta) and glial marker GFAP (cyan). (**C**) Representative whole organoid imaging of CUBIC-cleared brain organoids following GBM (white) entry into immature vs. more mature dorsal forebrain organoids. (**D**) UMAP of 19,954 individual malignant cells that passed QC and after exclusion of non-malignant cells based on copy number variations. Malignant cells were assigned to clusters based on Neftel *et. al.*, 2019^21^. (**E**) Proportion of cells across the six Neftel GBM cell types in immature and more mature dorsal forebrain organoids and in the parent tumor. (**F**) Two-dimensional representation of GBM states pre-engraftment vs. post-engraftment across each of the three independent GBM samples. (**G**) Area-proportional Venn diagram to visualize the overlap between differentially upregulated genes when GBM cells engrafted into immature and mature dorsal forebrain organoids were compared to the dissociated tumor pre-engraftment. Although many genes (187; select GO terms shown below the Venn diagram) were common across both microenvironments, a significant proportion of genes were also unique to each niche (134 genes in immature and 141 genes in mature microenvironments). (**H**) Volcano plot showing differentially upregulated genes after GBM engraftment in immature dorsal forebrain organoids (left) vs. mature dorsal forebrain organoids (right) and corresponding select GO terms shown below the volcano plot.

### The NPC-like state is reflective of neuronal gene signatures, with smaller contributions from OPC-like and radial glia-like programs

To increase the granularity of our cell type annotations and better define the lineage potential of NPC-like cells, we projected our data onto a reference GBM meta-atlas of single-cell transcriptomes from seven published GBM datasets, comprising 55 distinct tumors^40^. This analysis revealed that the NPC-like cell state in our data largely corresponded to a neuron-like transcriptional program in the reference atlas (**Fig. 4A-B**). Consistent with this observation, cell type proportion analyses based on reference mapping recapitulated the increase in neuron-like GBM cells post-engraftment, including the comparatively modest increase observed within mature organoids (**Fig. 4C**). Comparing the Neftel cell state annotations^21^ with those from the GBM meta-atlas^40^ further demonstrated that a vast majority of NPC-like cells align with the neuron-like program, with smaller subsets corresponding to OPC-like identities and a minor fraction with radial glia-like identities (**Fig. 4D**). Within organoids, GFP^+^ GBM cells indeed colocalized with the OPC-lineage marker, Olig2, and neuron marker, MAP2 (**Fig. 4E**).

**Fig. 4.**
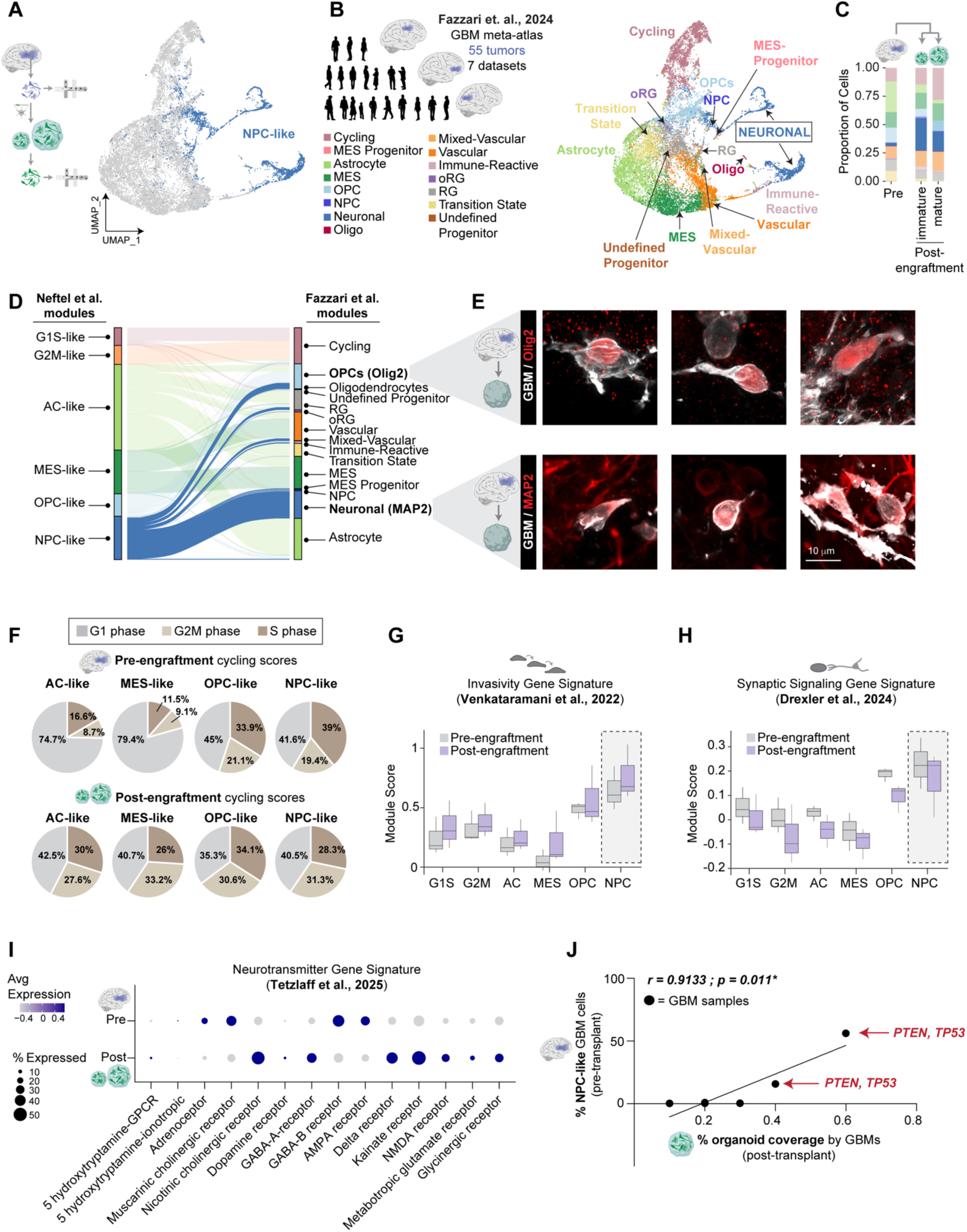
NPC-like GBM cells express neuron-like, invasion-related, and synaptic signaling-related genes post-engraftment. (**A-B**) Transcriptomic data from Fig. 3 (pre-engraftment vs. post-engraftment into immature vs. more mature organoids) projected onto a GBM meta-atlas from Fazzari *et. al.* 2024^40^ of single-cell transcriptomes from 55 patient tumors across 7 published datasets. (**C**) Proportion of cells across cell types from the GBM meta-atlas^40^ in immature and more mature dorsal forebrain organoids and in the parent tumor. (**D**) Sankey plot comparing the cluster annotations from Neftel *et. al.^21^* with the annotations from the Fazzari *et. al.* GBM meta-atlas. (**E**) Representative high-magnification images of GBM-engrafted organoids to show colocalization of direct-from-patient GBM cells (white) with the OPC lineage marker Olig2 (red) and the neuron lineage marker MAP2 (red). (**F**) Cell cycle scores from the Seurat pipeline were used to calculate the proportion of cells in G1, S, and G2M phases before and after engraftment, shown here based on Neftel *et. al.* cluster annotations of the transcriptomic data from Fig. 3. (**G**) Module scores for invasion-related genes^8^, displayed for GBM cells before vs. after organoid engraftment, shown here based on Neftel *et. al.* cluster annotations of transcriptomic data from Fig. 3. (**H**) Module scores for synaptic signaling-related genes^60^, displayed for GBM cells before vs. after organoid engraftment, shown here based on Neftel *et. al.* cluster annotations of transcriptomic data from Fig. 3. (**I**) Neurotransmitter receptor gene expression signatures^54^, displayed for GBM cells before vs. after organoid engraftment, using the transcriptomic data from Fig. 3. (**J**) Total proportion of cells assigned to the NPC-like state pre-engraftment for each of the *n=*6 GBM samples across Fig. 2-3 were correlated with the yield of GFP^+^ cells following FACS on dissociated, GBM-engrafted organoids (Spearman’s *r=*0.9133, *p=*0.011*). The two GBM samples with the greatest NPC-like cells pre-engraftment showed the highest yield of GFP^+^ cells post-engraftment and both these tumors showed genetic abnormalities in *PTEN* and *TP53*.

To investigate if there was any further molecular heterogeneity within the NPC-like state of GBM cells before and after engraftment, we subsetted and re-clustered NPC-like cells. We found an equal representation of the Neftel NPC1 subtype (includes OPC-related genes^21^) and NPC2 subtype (includes neuron-related genes^21^) (**Fig. S9A-C**). Following engraftment, we observed a modest increase in NPC1-like GBM cells relative to NPC2-like cells in immature microenvironments, potentially reflecting the activation of early neurogenic signaling programs (**Fig. S9C**). Notably, this NPC1-to-NPC2 ratio reverted to pre-engraftment levels within more mature organoids (**Fig. S9C**). We also sought to determine whether NPC-like cells were biased toward specific neuronal subtypes, such as excitatory or inhibitory neurons. We discovered that, upon engraftment, NPC-like GBM cells show a reduction in immature neuron-like signatures and an increase in both, inhibitory and excitatory neuron-like fate commitment (**Fig. S9D-F**). Thus, this shift may reflect changes in neurotransmitter signaling or receptor expression upon engraftment and entry of GBM cells into organoids, potentially driven by synaptic engagement between GBM and host organoid cells.

### NPC-like GBM cells are enriched in expression of invasion and synaptic signaling-related genes

We examined whether the post-engraftment increase in NPC-like GBM cells reflected an increase in proliferation or enhanced invasion of these cells within the organoid environment. We assessed this by scoring cell cycle phases and found that, prior to engraftment, NPC-like and OPC-like GBM cells showed more active proliferation (S-phase) than astrocyte-like and MES-like cells (**Fig. 4F**; top row). However, after engraftment, the distribution of cells across G1, S, and G2/M phases were largely comparable across all four major GBM cell types (**Fig. 4F**; bottom row). In contrast, NPC-like GBM cells continued to show greater expression of genes related to invasion^8^ and synaptic signaling^60^ after engraftment within the organoid microenvironment (**Fig. 4G, S9G**) Given that increased invasiveness may correlate with greater synaptic signaling, we next examined the landscape of neurotransmitter-related gene modules^54^ present on GBM cells (**Fig. 4H**). Compared to the dissociated, input tumor, GBM cells within organoids exhibited an upregulation in the expression of nicotinic cholinergic receptors, inhibitory GABA-A and glycinergic receptors, and several components of excitatory glutamatergic signaling, such as delta, kainate, NMDA, and metabotropic glutamate receptors (**Fig. 4I** and **Fig. S10A-B**). These findings are consistent with our observation that NPC-like cells post-engraftment acquire transcriptomic signatures resembling both excitatory and inhibitory neurons. Finally, to investigate the invasive potential of NPC-like GBM cells, we correlated the proportion of NPC-like cells *pre-engraftment* and the extent of organoid coverage by GBM cells *post-engraftment*, across all six direct-from-patient GBM samples used in this study (reflected in Fig. 2-3). We found that tumors with higher NPC-like composition prior to engraftment exhibited broader organoid coverage, suggesting that this cell type is indeed intrinsically more invasive or better equipped to adapt to the organoid niche (**Fig. 4I**). Interestingly, both NPC-rich GBMs in our cohort harbored mutations in *TP53* and *PTEN*. These data suggest potential shared intrinsic factors underlying the behaviors of this cell type within the neural microenvironment.

### GBM cells are more invasive in the dorsal forebrain microenvironment than the spinal cord

Although our data demonstrated that tumor cells converge onto a shared transcriptomic state across brain regions, we wanted to determine if these shared transcriptional profiles translate into similar infiltrative behaviors. We engrafted gliomasphere-derived cells for 14 days and found that they infiltrated forebrain organoids more robustly than spinal organoids [**Fig. 5A**; 316.1 μm (*p<*0.05) and 396.9 μm (*p<*0.01) in dorsal and ventral forebrain organoids, versus 196.4 μm in spinal cord], possibly reflecting the influence of distinct host cell compositions across these regions. Next, we exposed GBM-engrafted dorsal forebrain and spinal cord organoids to the nucleotide analog, EdU, throughout the 14-day engraftment period, followed by flow cytometry to quantify the total numbers of GFP^+^ tumor cells and the fraction of GFP^+^ tumor cells that are also EdU^+^ (the proportion of proliferating cells; **Fig. S11A**). Interestingly, there were more GFP^+^ tumor cells in the spinal cord than the dorsal forebrain (**Fig. S11B**), suggesting that the increased invasiveness of GBM in the dorsal forebrain was not attributable to greater overall tumor burden. Notably, the proportion of proliferating GBM cells was also higher in dorsal forebrain organoids (**Fig. S11C**). These findings suggest that the dorsal forebrain microenvironment, which is rich in excitatory neurons, supports a GBM cell state that is both highly invasive and proliferative.

**Fig. 5.**
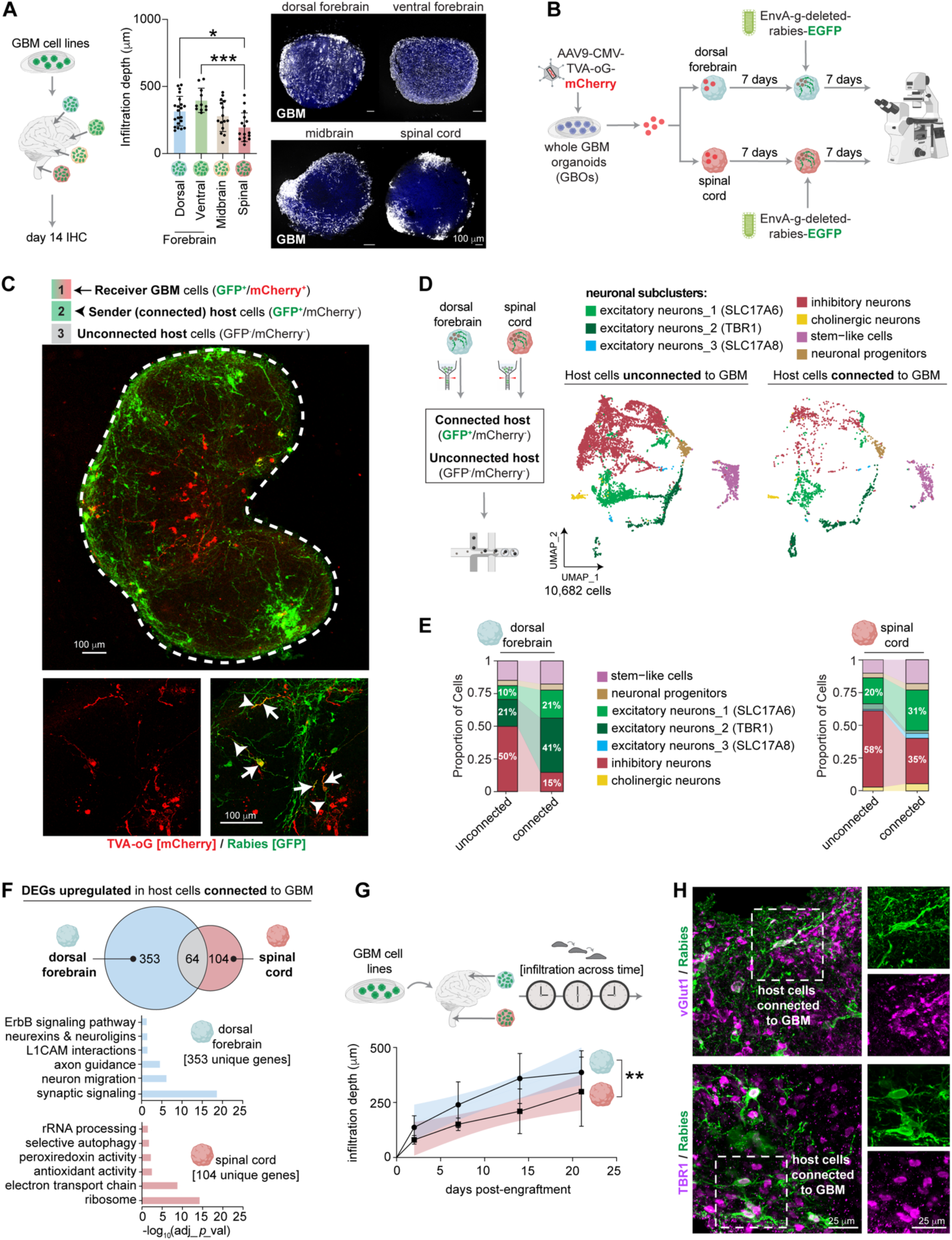
Host organoid cells synaptically connected to GBM cells are more likely to be TBR1^+^ excitatory neurons in the dorsal forebrain. (**A**) Infiltration depth of GBM cells from patient-derived cell lines engrafted for 14 days into dorsal forebrain, ventral forebrain, midbrain, and spinal cord organoids. Each dot is the GBM infiltration depth in a single organoid section. Data are from two independent GBM cell lines engrafted into two independent hiPSC-derived brain organoid lines (*n=*2 GBM lines x 2 organoid lines). Data were non-normal (based on Shapiro-Wilk) and analyzed by Kruskal-Wallis followed by the Dunn’s test to control for multiple comparisons. (**B**) Whole GBM organoids (GBOs) were transduced with AAV9-CMV-TVA-mCherry-2A-oG, following which GBM cells were engrafted into dorsal forebrain and spinal cord organoids. After 7 days, engrafted organoids were infected with EnvA G-deleted rabies-EGFP and 7 days later (14 days post-engraftment) organoids were fixed for immunohistochemistry. (**C**) Representative image of “receiver” GBM cells (arrow; GFP^+^ / mCherry^+^), synaptically connected “host” organoid cells (arrowhead; GFP^+^ / mCherry^-^), and synaptically unconnected “host” organoid cells (unmarked; GFP^-^ / mCherry^-^) at low and high magnifications. (**D**) Host organoid cells synaptically connected to GBM (GFP^+^ / mCherry^-^) and unconnected to GBM (GFP^-^ / mCherry^-^) were isolated by FACS and subject to single-cell captures. UMAP of 10,682 tumor-connected vs. tumor-unconnected host neuron-related subclusters. (**E**) Proportion of cells assigned to each of the neuron-related subclusters in the tumor-connected vs. tumor-unconnected host neurons in the dorsal forebrain and spinal cord niches. (**F**) Area-proportional Venn diagram to visualize the overlap between differentially upregulated genes when host cells that are synaptically connected to GBM are compared to the host cells synaptically unconnected to GBM in the dorsal forebrain (blue) vs. the spinal cord (red). GO terms related to genes that are uniquely upregulated (353 in the dorsal forebrain and 104 in the spinal cord) in the GBM-connected host fraction in each organoid niche are shown. (**G**) Infiltration depth of GBM cells from patient-derived cell lines engrafted for 2, 7, 14, and 21 days into dorsal forebrain and spinal cord organoids. Data are from two independent GBM cell lines engrafted into two independent hiPSC-derived brain organoid lines (*n=*2 GBM lines x 2 organoid lines). Each dot reflects the average infiltration depth for each individual “n”. Data were normally distributed (based on Shapiro-Wilk) and analyzed by 2-way ANOVA followed by Tukey; shown is the statistical main effect of region. (**H**) Representative image of tumor-connected host organoid cells expressing rabies (green) and colocalized with the excitatory neuron markers vGlut1 (magenta; top row) and TBR1 (magenta; bottom row) at low and high magnification. All error bars are mean + SD.

### Host organoid cells synaptically connected to GBM cells are enriched in TBR1^+^ excitatory neurons

Given greater infiltration and proliferation of GBM cells in the dorsal forebrain compared to spinal cord and our finding that GBM cells upregulate various excitatory neurotransmitter receptor modules after entry into organoids, we next asked whether these regional disparities could be explained by differences in synaptic connectivity between host and tumor. To investigate this, we used the pseudotyped rabies virus-based monosynaptic tracing technique (**Fig. 5B**). We exposed whole GBM organoids (GBOs) to AAV9-CMV-TVA-mCherry-2A-oG, allowing for expression of the TVA receptor and the rabies oG glycoprotein in tumor cells. We used GBOs for this experiment rather than direct-from-patient GBM cells, as TVA/oG expression required ∼3 weeks to peak and we wanted to avoid prolonged *in vitro* culture of freshly resected tumors. Following engraftment of the labeled tumor cells into brain organoids, exposure to EnvA G-deleted rabies-EGFP facilitates infection of TVA^+^ GBM cells with rabies, such that the GBM cells co-express GFP (from rabies) and mCherry (from TVA). Thus, the initial labeling of tumor cells with TVA limits the initial rabies infection to tumor cells alone, sparing host organoid cells. After the initial infection, trans-complementation by oG enables one round of retrograde transfer of rabies, so that host cells synaptically connected to GBM become labeled by GFP, but lack mCherry expression. Because host cells do not endogenously express oG, viral spread is restricted to first-order inputs. Using this method, we identified three cell populations: (1) “receiver” GBM cells (GFP^+^/mCherry^+^; arrow in **Fig. 5C**), (2) “sender” GBM-connected host organoid cells (GFP^+^/mCherry^-^; arrowhead in **Fig. 5C**), and (3) GBM-unconnected host organoid cells (GFP^-^/mCherry^-^; unlabeled cells in **Fig. 5C**). We validated the specificity of this system by confirming that rabies delivery after engraftment of *control* GBOs (without TVA label) yielded no GFP labeling (**Fig. S12A-B**).

We used FACS to separate GBM-connected host cells (GFP^+^/mCherry^-^) and GBM-unconnected host cells (GFP^-^/mCherry^-^) from dorsal forebrain and spinal cord organoids and performed single-cell RNA sequencing on each population (**Fig. S12C**). To ensure that our analyses were restricted to host-derived organoid cells, we excluded cells with detectable CNVs, validated by the expression of *XIST*, consistent with our use of a female GBO line engrafted into a male hiPSC-derived organoid (**Fig. S12D-E**). After confirming the purity of the dataset, we annotated the remaining non-malignant cell clusters and compared the compositions of GBM-connected and unconnected host populations. Across both regions, GBM-connected host cells were less likely to be inhibitory neurons (**Fig. S12F-H**). In contrast, excitatory neurons were enriched among GBM-connected cells in the dorsal forebrain but were reduced in the spinal cord (**Fig. S12G**), indicating regional biases in neuronal subtypes engaged by GBM cells. Although there were also modest increases in OPCs, oligodendrocytes, and radial glia/astrocytes in GBM-connected host cells (**Fig. S12G**), we focused our analysis into the neuronal compartment.

After re-clustering neuronal cells (**Fig. 5D; Fig. S13A-C**), we found that host neurons expressing the excitatory transporter gene, *SLC17A6* [vGlut2], were connected to GBM in both the dorsal forebrain and the spinal cord, indicating a shared preference for this glutamatergic input in both regions. However, the dominant neuron subtype connected to GBM in the dorsal forebrain co-expressed deep-layer marker, TBR1 and the cortical excitatory marker *SLC17A7* [vGlut1] (**Fig. 5E**; 41% of total neurons in the GBM-connected fraction in the dorsal forebrain versus 21% in the GBM-unconnected fraction). This suggests that, within the dorsal forebrain, GBM has a greater likelihood of receiving synaptic input from deep-layer excitatory neurons. In contrast, inhibitory neurons dominated the GBM-connected spinal cord organoids (35% of total neurons in this fraction; **Fig. 5E**), even though they made up a smaller proportion of the total neuron population when compared to the unconnected fraction in the same region (58%).

On comparing DEGs between tumor-connected and unconnected neurons, we found greater numbers of transcriptional changes in the dorsal forebrain vs. spinal cord (upregulated DEGs in **Fig. 5F**; downregulated DEGs in **Fig. S13D-F**; **Supplementary Table 6**), indicating more extensive transcriptomic remodeling following the establishment of host-GBM synapses in the forebrain. GO analyses revealed that GBM-connected dorsal forebrain neurons were enriched for pathways related to axon guidance, neuron migration, synaptic signaling, ErbB signaling, and neurexins-neuroligins. GBM-connected spinal neurons upregulated genes linked to stress-response and mitochondrial bioenergetics (**Fig. 5F**).

Consistent with GO terms related to migration and axon guidance, we found that GBM cells were indeed more migratory in the dorsal forebrain versus the spinal cord across days 2-21 post-engraftment (statistical main effect of *region*; *p<*0.01; **Fig. 5G**). In addition, we validated that GBM-connected host cells in the dorsal forebrain co-localized with excitatory cortical neuron marker vGlut1 and the deep-layer, long-range projection neuron marker TBR1 (**Fig. 5H**). Together, these findings suggest that GBM engages deep-layer TBR1^+^ excitatory neurons in the dorsal forebrain and this neuron-tumor crosstalk may underlie the enhanced invasiveness of GBM in this region.

## DISCUSSION

A central question we sought to address is whether GBM cells adopt unique or universal gene expression signatures across the distinct spatiotemporal environments of the nervous system. We used a fully human platform, engrafting cells from the same GBM samples into dorsal forebrain, ventral forebrain, midbrain, and spinal cord organoids, derived in parallel from the same iPSC lines and differentiations. Although organoids are inherently reductionist and lack immune cells or an endogenous vasculature, this study was designed to dissect how specific regional host neural cells interact with invading GBM cells in a manner that is not readily feasible in an *in vivo* setting. We found that, after engraftment, GBM tumors are depleted in astrocyte-like transcriptional identities and enriched in NPC-like identities, irrespective of brain region or the maturational ages of the organoid. NPC-like GBM cells expressed neuronal genes, including those related to excitatory and inhibitory neuronal types, and GBM cells upregulated multiple neurotransmitter-related receptors following engraftment. Notably, NPC-like GBM cells maintained higher expression of invasion-related genes within the organoid niche, consistent with prior work showing that deeply infiltrative gliomasphere cells express neuronal markers, such as TUJ1^61^. We further observed that tumors enriched in NPCs before engraftment correlated with broader organoid coverage, underscoring the translational potential of targeting this specific infiltrative GBM cell type. These findings align with recent studies showing that neuron-like GBM cells are enriched at recurrence^62-64^, supporting the idea that these cells may promote GBM progression. GBM-engrafted organoids are thus an ideal testbed for evaluating whether therapeutic agents selectively targeting NPC/neuron-like GBM cells impede tumor infiltration. This is noteworthy since these specific cell types are not fully captured in gliomasphere cell lines^36^, perhaps because of selection bias^65^ or because these cells may be difficult to retain after enzymatic dissociation of the parent tumor^22^.

We compared our post-engraftment transcriptomic data to the matched, dissociated input GBM to control for patient-specific genetic variations. Although we are low in statistical power, tumors rich in NPCs harbored mutations in *PTEN and TP53*, suggesting a potential role for tumor-intrinsic factors in driving transcriptomic changes following organoid engraftment. In support of this idea, rodent models show that dual, but not singular deletion of *PTEN* and *TP53* promote tumorigenesis, potentially via increased c-Myc activity^66^. We restricted our analyses to IDH1 wild type GBM tumors, which we previously reported to be enriched in astrocytes stalled at an intermediate stage of maturation^50^. If this stage is a checkpoint in glial fate determination, it is possible that our findings also reflect a plastic cell state that is easily swayed towards an NPC/neuron-like fate following entry of tumor cells into the neural niche and reception of pro-neurogenic cues. It will be critical to define if IDH1 mutant gliomas, with their more mature astrocytic cells^50^, more stable differentiation trajectories^67^, milder pathology^68-71^, and improved patient prognoses^72-74^ also similarly display greater NPC/neuron-like cell states following entry into the neural microenvironment.

Despite a shared transcriptomic state across regions, GBM cells were more infiltrative in dorsal forebrain as compared to spinal cord organoids. Interestingly, GBM cells were also more proliferative in the dorsal forebrain, supporting the recently reported “go-AND-grow” characteristics of GBM cells during initial stages of brain colonization^75^. We propose that the dorsal forebrain microenvironment promotes a more disperse and mitotically active phenotype in GBM cells, one that is potentially capable of escaping resection and seeding new lesions through simultaneous invasion and proliferation. In contrast, the spinal cord may support an interconnected tumor network that is spatially confined and functionally dormant. Whether GBM cells in the dorsal forebrain eventually transition to a more latent and interconnected state remains an open question. Our findings also suggest that GBM invasiveness depends not only on crosstalk with host neurons^8,76^, but also on the specific neuronal subtypes and regional identities that tumor cells engage with.

We found that inter-regional differences in GBM infiltration were not explained by differences in total tumor burden. Given emerging evidence that invasive gene signatures in GBM may correlate with synaptic signaling^54^, we used pseudotyped rabies virus-based monosynaptic tracing to map the landscape of host neural cells that synapse onto GBM in the dorsal forebrain versus spinal cord. We made two major observations: (1) vGlut2-expressing excitatory neurons are more likely to synapse onto GBM cells in both regions, whereas inhibitory neurons are less likely to be synaptic partners of GBM, and (2) a subtype of excitatory neurons that expresses the deep-layer neuron marker TBR1 is the predominant synaptic partner of GBM in the dorsal forebrain. We speculate that TBR1^+^ neurons may release paracrine cues that promote GBM invasion (and proliferation) in the dorsal forebrain. In the spinal cord, inhibitory interneurons were the primary senders of synaptic input to GBM and may release molecules that impede GBM infiltration in this region.

Whether GBM cells also engage superficial layer neurons in more mature forebrain organoids is to be investigated since this population was less abundant at the age when our rabies tracing was performed. Notably, our data revealed that TBR1^+^ forebrain neurons preferentially synapse onto GBM and co-express SLC17A7, a marker for long-range projection neurons in the cortex^77^. Since TBR1^+^ neurons are involved in circuits connecting the cortex to thalamic and subcortical areas^78^, our findings suggest that GBM cells preferentially engage long-range excitatory pathways—even in the absence of fully intact circuits, as in our brain organoid models. Future studies in assembloid models that recapitulate inter-regional interactions could be useful to clarify if long-range excitatory projections from TBR1 or SLC17A7-expressing neurons in the forebrain can also modulate tumor infiltration and synaptic integration in distal areas. These long-range circuits may grant GBM cells access to a diverse pool of neuromodulatory signals^4,53^ by expanding the number and range of synaptic inputs, and may potentially accelerate GBM infiltration and integration, ultimately contributing to the aggressive nature of this tumor. Examining this in future work will have notable clinical implications, as GBM tumors invade into hindbrain structures at late-stage disease^55^. Collectively, our data suggest that targeting brain region-specific or circuit-specific tumor-neuron interactions may be functionally important and therapeutically targetable in glioblastoma.

## METHODS

### Generation of human iPSC-derived forebrain, midbrain, and spinal cord organoids

One male (8858.3) and one female (C4.1) hiPSC line were used throughout the experiments in this manuscript. These lines were sourced from the Pasca lab at Stanford University (8858.3) or Coyne Scientific (C4.1) and were generated from unidentified healthy individuals. hiPSC lines were regularly tested for mycoplasma contamination and genomic integrity via SNP array. hiPSC lines were cultured on a vitronectin (ThermoFisher, A14700)-coated plate in Essential 8 medium (ThermoFisher, A1517001) and passaged in clumps with EDTA or ReleSR (STEMCELL Technologies, 05872) for maintenance, expansion, and freezing. For forming organoids, hiPSC colonies at 80-90% confluency were detached using Accutase (VWR, 10761-312) and formed into 3D embryoid bodies (EBs) overnight in AggreWell 800 plates (STEMCELL Technologies, 34811) at 3 million cells per well in Essential 8 medium supplemented with 10μM rock inhibitor Y-267632 (Tocris, 1254). The next day, EBs were transferred to 100mm dishes that were made ultra-low attachment with Anti-Adherence Rinsing Solution (STEMCELL Technologies, 07010). EBs were patterned to dorsal forebrain, ventral forebrain, midbrain, and spinal cord identities using previously established protocols and patterning recipes^44-48^.

For forebrain, EBs were cultured in Essential 6 medium (ThermoFisher, A1516401) supplemented with Penicillin/Streptomycin (VWR, 16777-164) from days 1-5, and then switched to Neurobasal-A (US Bio, N1020-02) medium supplemented with B-27 (ThermoFisher 12587010), Glutamax (ThermoFisher, 35050061), and Penicillin/Streptomycin. To pattern EBs to dorsal fates^44^, dual SMAD inhibition was achieved using Dorsomorphin (Sigma-Aldrich, P5499; 2.5μM) and SB-431542 (Selleck Chemicals, S1067; 10μM) until day 5, following which EGF (R&D Systems, 236-EG; 20ng/mL) and FGF2 (R&D Systems, 233-FB; 20ng/mL) were added until day 25, and then BDNF (PeproTech, 450-02; 20ng/mL) and NT-3 (PeproTech, 267-N3-005; 20ng/mL) were added until day 44. From day 45 onward and until termination, dorsal forebrain organoids were cultured only in Neurobasal-A medium supplemented with B-27, Glutamax, and Penicillin/Streptomycin. To pattern EBs to ventral fates^46^, dual SMAD inhibition was achieved using 2.5μM of Dorsomorphin and 10μM SB-431542 until day 5. Wnt pathway inhibitor XAV939 (STEMCELL Technologies, 72674; 2.5μM) was added between days 3-5. Next, organoids were cultured in 20ng/mL EGF, 20ng/mL FGF2, and the Wnt inhibitor IWP-2 (Selleck Chemicals, S7085; 2.5μM) until day 24, with the smoothened agonist SAG (Sigma-Aldrich, 566660; 100nM) added between days 11-24 for ventralization. From day 25 to 44, organoids received 20 ng/mL BDNF and 20ng/mL NT-3 and then cultured only in Neurobasal-A with B-27, Glutamax, and Penicillin/Streptomycin, until termination.

For midbrain^47^, EBs were cultured in midbrain induction medium (MIM), containing 50:50 DMEM/F12 (ThermoFisher, 11320-033) and Neurobasal-A, supplemented with N-2 (ThermoFisher, 17502048), MEM-NEAA (ThermoFisher, 11140050), 2-mercaptoethanol (100mM), B27, Glutamax, and Penicillin/Streptomycin until day 7. From day 8, they were cultured in midbrain differentiation medium (MDM), containing Neurobasal-A supplemented with N-2, MEM-NEAA, 2-mercaptoethanol, B27, Glutamax, and Penicillin/Streptomycin. To pattern EBs to midbrain fates, heparin (Sigma-Aldrich, H3149-25KU; 1μg/mL), SB-431542 (10μM), Noggin (PeproTech, 120-10C; 200ng/mL), and Wnt activator CHIR99021 (Selleck Chemicals, S1263; 0.8μM) were added until day 7. Sonic hedgehog (Shh; PeproTech, 100-45; 100ng/mL) and FGF8b (PeproTech, 100-25; 100ng/mL) were added between days 4-7 for patterning to mesencephalic fate. From day 8 onward and until termination, the midbrain organoids received BDNF (10ng/mL), GDNF (PeproTech, 450-10; 10ng/mL), cAMP (Sigma-Aldrich, D0627; 125μM), and L-ascorbic acid (Fisher Scientific, 50-990-141; 100μM).

For spinal cord^45^, EBs were cultured in Essential 6 medium supplemented with Penicillin/Streptomycin until day 4 and then switched to Neurobasal-A medium supplemented with B-27, Glutamax, and Penicillin/Streptomycin. Dual SMAD inhibition using both Dorsopmorphin (2.5μM) and SB-431542 (10μM) was performed until day 4, with the Wnt activator CHIR99021 (3μM) added between days 3-4. From days 5-16, the growth factors EGF (20ng/mL) and FGF2 (20ng/mL) were added, in addition to Wnt activator CHIR99021 (3μM) and the hindbrain patterning agent retinoic acid (Sigma-Aldrich, R2625; 0.1μM). The smoothened agonist SAG (100nM) was added between days 9-16. From days 17-22, spinal cord organoids were cultured in N-2 supplement, BDNF (20ng/mL), cAMP (50 μM), L-ascorbic acid (200 μM), insulin-like growth factor I (IGF-I; Peprotech, 100-11; 100ng/mL), and Notch pathway modulator DAPT (STEMCELL Technologies, 72082; 2.5 μM). From day 22 onward and until termination, spinal cord organoids received N-2 supplement, BDNF (20ng/mL), cAMP (50 μM), L-ascorbic acid (200 μM), and IGF-I (100ng/mL).

Region-specific brain organoids patterned using these recipes were regularly validated via bulk RNA-sequencing and immunohistochemistry for canonical markers of regionalization.

### GBM tissue procurement

Fresh surgically resected GBM tissues were obtained from consenting and de-identified individuals, following diagnostic genomic profiling, and in full compliance with policies outlined by the Emory School of Medicine Institutional Review Board under the protocol IRB00045732. We used primary (non-recurrent) IDH1 wildtype GBM tissues in this study (Supplementary Table 1). Safe surgical resection of the GBM was performed by the operating neurosurgeon using direct vision, imaging, and fluorescence. GBM tissues were immediately collected in ice-cold Hibernate-A medium (ThermoFisher, A1247501), as described in our prior work^50^, and used for dissociations or minced to form whole GBM organoids within 1 hour following resection.

### Establishment of direct-from-patient GBM cultures, GBM cell lines, and whole GBM organoids

For the direct-from-patient cultures, tumor tissues were dissociated based on our previously established protocol^50^. In brief, pieces with extensive necrosis or hemorrhage were first removed from the tissue block using forceps and then the tumor tissues were chopped with a scalpel and incubated in papain (Worthington Biochemical, LS003126; 45 units/mL) supplemented with DNAse at 34 °C for 1 hour. After papain was quenched with the protease inhibitor ovomucoid (Worthington Biochemical, LS003086), we performed gentle mechanical trituration to obtain a single-cell suspension, and then filtered it through a 40 μm strainer. Following cell counts, we used this cell suspension immediately for single-cell captures, viral labeling, and engraftment. For single-cell captures, GBM cell pellets were resuspended in a buffer containing DPBS without calcium/magnesium (VWR, 45000-434), 1mM EDTA (ThermoFisher, 15575020), 25mM HEPES pH 7.0 (Sigma-Aldrich, 83264), and 1% heat-inactivated FBS (R&D Systems, S11150H). For viral labeling, GBM pellets were resuspended in NeuroCult NS-A medium (STEMCELL Technologies, 05750) supplemented with N2, B27, sodium pyruvate (ThermoFisher, 11360070; 1mM), Penicillin/Streptomycin, and L-glutamine (ThermoFisher, 25030081; 2mM).

GBM cell line spheres were established based on previously published protocols^79,80^ and ∼1 million cells were maintained in 100mm dishes in DMEM/F12+Glutamax (ThermoFisher, 10565018), supplemented with B27 (Invitrogen, 17504044), Penicillin/Streptomycin, heparin (5 μg/mL), EGF (20ng/mL), and FGF2 (20ng/mL). GBM spheres were enzymatically passaged as necessary for maintenance or expansion using Accutase (VWR, 10761-312) and frozen in BAMBANKER (VWR, 101974-112).

GBM organoids (GBOs) were prepared using published protocols^26,27^. In brief, after excluding necrotic and hemorrhaging bits, tumor tissues were washed with ice-cold DPBS (ThermoFisher, 14040133) and carefully cut into 0.5-1mm chunks using forceps and fine-spring dissection scissors (Fine Science Tools, 15750-11) in ice-cold GBM dissection medium, which contains Hibernate-A supplemented with Glutamax and antibiotic-antimycotic (ThermoFisher, 15240062). GBM tissue pieces were washed with room temperature DPBS, incubated in RBC lysis buffer (ThermoFisher, 00433357) for 10 minutes at room temperature, washed with DMEM/F12, and then maintained in ultra-low attachment plates in GBO medium, containing 50:50 DMEM/F12 and Neurobasal (ThermoFisher, 21103049) media supplemented with MEM-NEAA, Glutamax, N2, B27, Penicillin/Streptomycin, insulin (Sigma-Aldrich, 19278), and 2-mercaptoethanol (ThermoFisher, 21985023). GBOs round up within 1-2 weeks after formation, can be passaged when they reach 1-2mm in diameter by sequential bisection, and cryopreserved^26,27^.

### Viral labeling of GBM cells

GBM cells were labeled with pLenti-CMV-GFP (Addgene, 17446-LV), regardless of source (direct-from-patient vs. cell lines vs. GBOs), at a multiplicity of infection (MOI) of 5. For the physical labeling, direct-from-patient GBM cells in suspension were incubated with the lentivirus in a tube at 37 °C and 5% CO_2_ for 4 hours on a shaker, set at 150rpm. After 4 hours, cells were washed two times with pre-warmed media before counting and organoid engraftment. To optimize this protocol, initial experiments also compared this method of GFP labeling in suspension to overnight (∼18 hours) GFP labeling in suspension and to cells plated on a Matrigel substrate (VWR, BD354277) and exposed to the GFP lentivirus 1 hour following plating, for total of 24 hours. This latter method of pre-plating on Matrigel involved gently passaging cells with Accutase following the 24 hours of GFP labeling, cell counts, and subsequent organoid engraftment. Based on our characterization of these culture and lentiviral exposure conditions, we proceeded with the 4-hour suspension labeling of direct-from-patient GBM cells for all subsequent experiments described in this manuscript.

For GBM cell lines, spheres passaged by Accutase into a single-cell suspension were exposed to the same GFP lentivirus as described above at MOI 5 for 48 hours, following which cells were washed with media, and the spheres were allowed to reform. These GFP^+^ GBM cell line spheres were then maintained, passaged, and cryopreserved as per the regular, established protocols.

To label whole GBM organoids, GBOs were subject to papain-based dissociation, as described above for direct-from-patient GBM cultures, except that the concentration of papain was 37.5 U/mL and we reduced the number of wash steps and completely excluded the filtering step. After the GBOs were dissociated, we performed cell counts and proceeded with the same 4-hour suspension labeling of cells with the GFP lentivirus, as described above.

### GBM cell engraftment into brain organoids

We initially attempted multiple physical engraftment strategies, such as co-culturing brain organoids with GFP-labeled GBM cells in v-bottom or flat-bottom 96-well plates, in 24-well plates left tilted in an incubator, or via a hanging drop method. However, we noticed that maintaining GFP-labeled GBM cells in suspension on top of a brain organoid in a v-bottom (conical) low-protein binding Eppendorf tube gave us the most reproducible success of engraftment across multiple distinct GBM samples, including direct-from-patient GBM cells, accutase-passaged GBM cell line spheres, and dissociated GBOs. We engrafted 50,000-100,000 GBM cells per organoid, maintained in 500μL of region-specific organoid media for ∼36-48 hours. Next, the un-engrafted GBM cells were washed with three media exchanges, and GBM-engrafted organoids were transferred from Eppendorf tubes into ultra-low attachment plates. GBM-engrafted organoids were maintained in region-specific organoid media until the end of the experiments. To minimize batch and technical variability, we engrafted the same GBM sample into forebrain, midbrain, and spinal cord organoids derived in parallel from the same hiPSC lines and differentiations. In a similar vein, we also engrafted the same GBM sample into immature vs. more mature dorsal forebrain organoids derived from the same hiPSC lines. For experiments comparing distinct regional environments, the ages of region-specific brain organoids were between days 38-59 at engraftment. For experiments comparing engraftments across developmental ages, we used dorsal forebrain organoids between days 36-64 to mimic younger (more immature) microenvironments and days 208-279 for to mimic older (more mature) microenvironments. These organoid ages were selected based on their availability in our Brain Organoid Hub at the time of GBM tissue harvest and to ensure sufficient temporal and maturational separation between immature and mature dorsal forebrain organoids. This separation aligns with the neurogenic-to-gliogenic switch that we have previously characterized in this model^50,51^.

### Dissociation of GBM-engrafted organoids and FACS sorting

At the end of the engraftment period, GBM-engrafted organoids were dissociated into a single-cell suspension, using the same papain-based method described above for dissociating fresh surgical GBM resections. However, we modified the concentration of papain based on the organoid age [7.5 U/mL for younger (immature) organoids and 25 U/mL for older (mature) organoids], reduced the number of wash steps, and excluded the filtering step. The single-cell suspension was pelleted and resuspended in FACS buffer, containing DPBS without calcium/magnesium, supplemented with 1mM EDTA, 25mM HEPES pH 7.0, and 1% heat-inactivated FBS. We maintained unstained and single-stained samples as controls for fluorescence compensation and establishing gates during the FACS sorting procedure. FACS was performed on a BD Biosciences FACSAria II Cell Sorter or the BD FACSDiscover S8 Cell Sorter at the Emory Flow Cytometry Core. Following FACS, the GFP^+^ GBM cells and GFP^-^ host cells were either immediately captured for single-cell RNA sequencing or fixed in paraformaldehyde and frozen for latter single-cell captures using the multiplexed FLEX chemistry from 10x genomics.

### EdU exposures

GBM-engrafted organoids were fed with organoid media containing the thymidine analog 5-Ethynyl-2-deoxyuridine (EdU; 10μM) throughout the 14-day engraftment period, with media exchanges three times per week. After 14 days, GBM-engrafted organoids were dissociated as described above and the single-cell suspension was fixed and stained for EdU with the Click-IT EdU Alexa Fluor 647 Flow Cytometry Assay Kit (ThermoFisher, C10424), as per the manufacturer’s instructions. As controls for the flow cytometric assessment of proliferation indices, we used unstained and single-stained samples.

### Immunohistochemistry

Organoids were fixed with 4% paraformaldehyde (Electron Microscopy Sciences, 15710) for 3-4 hours at 4 °C and then transferred to 30% sucrose for 24-48 hours. After the organoids sank, they were thoroughly washed in optimal cutting temperature (OCT) compound (VWR, 25608-930), embedded in cryomolds in OCT, and stored at -80 °C. Cryomolds were cryosectioned at 70 μm thickness, transferred onto glass slides (ThermoFisher, 1250015), and stored at -80 °C. Cryosectioned organoids were washed with PBS containing 0.01% Triton X-100 (Sigma-Aldrich, X100) three times to remove OCT and blocked in PBS containing 10% normal donkey serum (Jackson ImmunoResearch, 0170001219) and 0.3% Triton X-100 for 60 minutes at room temperature. Sections were then incubated overnight at 4 °C in the same blocking buffer, supplemented with the following primary antibodies: Chk anti-GFP (Aves, GFP1020; 1:1500), Ms anti-Ki67 (BD Biosciences, B550609; 1:50), Rb anti-Sox2 (Cell Signaling Technology, 3579S; 1:500), Rb anti-GFAP (Agilent Dako, Z0334; 1:1500), Rb anti-Pax6 (Cell Signaling Technology, 60433S; 1:100), Ms anti-Nestin (Abcam, ab22035; 1:500), Ms anti-NKX2.1 (Santa Cruz Biotechnology, sc-53136; 1:200), Rb anti-GABA (Sigma-Aldrich, A2052; 1:500), Ms anti-Tyrosine Hydroxylase (ThermoFisher, MA1-24564; 1:100), Ms anti-ISL1 (DSHB, 39.4D5; 1:200), Rb anti-VAChT (Synaptic Systems, 139103; 1:300), Ms anti-MAP2 (Abcam, ab11267; 1:1000), Rb anti-Olig2 (Millipore, AB9610; 1:200), Rb anti-mCherry (Abcam, ab167453; 1:500), Rb anti-TBR1 (Abcam, ab31940; 1:500), and Rb anti-vGlut1 (ThermoFisher, 482400; 1:500). The next day, the sections were washed three times with PBS containing 0.01% Triton X-100 and incubated with the nuclear agent Hoechst 33342 (ThermoFisher, H3570; 1:2000) and secondary antibodies in the same buffer as the primary antibodies, for 1 hour at room temperature protected from light. Unbound secondary antibodies were washed off with three PBS exchanges and slides were coverslipped using Fluoromount-G mounting medium (ThermoFisher, 00495802). Slides were air dried and then imaged on a confocal microscope (Leica Stellaris 5; Las-X).

For comparison, groups were always imaged at the same laser gain and intensity settings, as well as the same zoom, line average, and resolution settings. Images were always coded and investigators blinded to the groups quantified the infiltration of GBM cells within organoid sections. We relied on two methods: (1) infiltration depths of individual GBM cells within the organoid, quantified by measuring the shortest distance between the rim of the organoid section and 5 randomly selected GBM cells in the central core of the organoid section, and (2) area of the organoid section that is covered by GBM cells after binning each organoid into 4 equally divided portions and assessing area in the outer rim (*i.e.,* in outer 25% of the organoid) and inner core (*i.e.,* in the inner 25% of the organoid).

### Tissue clearing

We used CUBIC to optically clear GBM-engrafted organoids^81-83^. In brief, organoids were fixed in 4% paraformaldehyde and 4% sucrose at 37 °C for 20 minutes and then 4 °C overnight, after which organoids were washed with PBS and incubated in CUBIC-L solution (TCI Chemicals, T3740) at 37 °C. After 48 hours, organoids were washed with PBS and stained with the primary antibody Chk anti-GFP (Aves, GFP1020; 1:1500) in PBS that contains 10% normal donkey serum and 0.2% Triton X-100 at 37 °C for 48 hours. Following PBS washes to remove unbound primary antibody, organoids were incubated in secondary antibody in PBS containing 10% normal donkey serum and 0.2% Triton X-100 at 37 °C for 48 hours and protected from light. Organoids were washed with PBS containing 0.1% Triton X-100 and transferred onto a glass-bottom 96-well plate (Corning, 4580) in CUBIC-R solution (TCI Chemicals, T3741) for at least 48 hours. Cleared organoids were then imaged on a confocal (Leica) microscope and z-stack images were collapsed into a maximum projection image to visualize total coverage of organoids by engrafted GFP^+^ GBM cells.

### Pseudotyped rabies virus-based monosynaptic tracing

To examine host-GBM synaptic connectivity, we used the pseudotyped rabies virus-based monosynaptic tracing technique, which has recently been used in other GBM models^52-54^ and has also been applied to hiPSC-derived brain organoids^45^. In brief, we transduced GBOs with AAV9-CMV-TVA-mCherry-2A-oG (Salk Institute) at 1:10 dilution. This construct allows for expression of the TVA receptor and the oG glycoprotein. We waited ∼3 weeks to allow the mCherry expression to peak, following which we dissociated GBOs and engrafted GBM cells into brain organoids, as per the method described above. After 7 days, we infected GBM-engrafted organoids with rabies (EnvA-g-deleted rabies-EGFP) at 1:100 dilution. The TVA receptor, expressed only on GBM cells, allows for the selective entry of the rabies into the GBM cells. Trans-complementation with the oG glycoprotein then promotes one round of retrograde transfer of the rabies into presynaptic partners (*i.e.,* host neural cells) of GBM cells. Using this method, we were able to detect receiver cells (GBM), labeled GFP^+^/mCherry^+^, synaptically connected host cells (GFP^+^/mCherry^-^), and synaptically unconnected host cells (GFP^-^/mCherry^-^). After 7 days (*i.e.,* 14 days post-engraftment), we either formalin fixed organoids for immunostaining or dissociated organoids for FACS sorting to separate synaptically connected and synaptically unconnected host cells for single-cell captures.

### Bulk RNA sequencing

Total RNA was extracted using the RNeasy Micro Kit (Qiagen, 74004), according to the manufacturer’s instructions. Bulk RNA libraries were prepared with NEBNext Ultra II RNA kit using polyA selection and sequenced on an Illumina 2x150 bp platform through Admera Health, at a targeted depth of 40 million paired end reads (20 million in each direction) for each sample. Fastq files were trimmed using Trimmomatic^84^, reads were mapped onto the hg38 reference using STAR aligner^85^, and the featureCounts software^86^ was used for read summarization to generate raw count matrices. Counts were normalized using TMM from edgeR^87^.

### Single-cell RNA sequencing

Single-cell captures and library preparation were performed on the chromium platform from 10x genomics, following the manufacturer’s instructions. We used either the Chromium Next-GEM single cell 3’ Kit v3.1 (10x genomics, 1000268) or Chromium GEM-X single cell 3’ Kit v4 (10x genomics, 1000691) for paired single-cell RNA sequencing comparing pre-engraftment transcriptomes to post-engraftment transcriptomes when using direct-from-patient GBM samples. For initial optimization of lentiviral exposure/culture conditions and for all paired single-cell RNA sequencing using GBM cell line spheres, we used the Chromium Next GEM Single Cell Fixed RNA kit (using the 10x genomics FLEX chemistry; 1000476), which allowed us to formalin fix, freeze, and ultimately multiplex samples for parallel captures. However, since this platform tends to elicit loss of cells during processing and hybridization, we only used the FLEX chemistry for samples where the yield following FACS was high. For rabies-based experiments, we used the Chromium GEM-X on-chip multiplexing (OCM) 3’ Chip Kit v4 4-plex (10x genomics, 1000779). Libraries were assessed on a Bioanalyzer (Agilent, 5067-4626) and sequenced at a target depth of at least 50,000 reads per cell on an Illumina 2x150 bp platform through Admera Health. Fastq files were processed using the 10x genomics cellranger count or cellranger multi pipelines with default parameters and aligned to the hg38 reference.

Additional data processing was completed on R Studio following the standard Seurat pipeline. For quality control, cells with >20% mitochondrial genes were excluded. The global-scaling normalization method “LogNormalize” was used and cells were clustered in the PCA space using FindNeighbors and FindClusters functions. Feature plots, dot plots, and differential gene expression analyses were performed on the raw RNA counts and according to the default Seurat parameters. Clusters were annotated either manually, based on the expression of canonical cell type-specific markers, or computationally, based on the highest module score calculated using Seurat’s AddModuleScore() function, incorporating the top genes for each GBM cell type that was defined by Neftel *et. al.,* 2019^21^. In addition, cluster annotations using the Neftel *et. al.* method were also projected onto a reference GBM meta-atlas^40^. Gene ontology (GO) analyses were performed using gProfiler^88^. Gene set enrichment analyses (GSEA) were performed using clusterProfiler^89^ and the Molecular Signature Database (MSigDB) with Hallmark (H) or curated (C5:GO) gene sets. Area-proportional venn diagrams were created using the eulerr package^90^. Sankey plots were created using the ggalluival package^91^. Invasion-related^8^, synaptic signaling-related^60^, and neurotransmitter receptor-related^54^ gene signatures were obtained from prior work, and module scores were calculated using Seurat’s AddModuleScore() function. CellChat^57^ analyses were performed to obtain ligand-receptor relationships in GBM cell line-engrafted brain organoids.

### Copy number variation (CNV) analyses to identify malignant vs. non-malignant GBM cells

We profiled for copy number variations (CNVs) using the inferCNV package^92^ with default parameters. For direct-from-patient GBM samples, immune cell clusters served as our “reference”. We quantified CNV correlation coefficients to separate malignant vs. non-malignant cells, as described in our prior work^50^. In short, we correlated the CNV signal of each cell with the average CNV signal across all cells from the same tissue. For post-engraftment samples, we correlated the CNV signal with the mean CNV signal across all cells of the matched pre-engraftment tissue to account for tumor sample-intrinsic CNV patterns. Where possible, this pipeline was validated by visualizing the expression of *EGFP* post-engraftment or, in cases where we were able to engraft GBM into organoids of the opposite sex, we validated our CNV pipeline by visualizing the expression of Y-chromosome genes *DDX3Y*, *USP9Y, KDM5D,* and *UTY*, or the expression of *XIST*. In this manner, we identified malignant vs. non-malignant cells, with the latter potentially introduced in our data as false positives from the FACS procedure or infiltrating cells from the tumor margin that were included in the resection.

### Statistical testing and reproducibility

Statistical analyses were performed in GraphPad Prism or using R Studio. We confirmed normality of data distribution using Shapiro-Wilk and then either employed appropriate ANOVAs (based on the number of independent variables; normally distributed data) or Kruskal-Wallis followed by the Dunn’s test to control for multiple comparisons (non-normal data). For two-sample comparisons, we employed the unpaired two-tailed *t* test (normally distributed data).

### Data and code availability

Sequencing files (Fastq, bam, featurecounts, etc.) will be made available through the Gene Expression Omnibus (GEO). Accession codes will be available before publication. R code for analysis will be made available on the Sloan Lab github (https://github.com/sloanlab-emory) before publication.

## ACKNOWLEDGMENTS

This study was supported by NIMH R01 MH125956, NINDS R01 NS123562, and the Sontag Foundation Distinguished Scholars Award (S.A.S.), as well as the American Cancer Society Postdoctoral Fellowship PF-24-1243136-01-CCB (T.N.B.) and a Postdoctoral Scholar Award from Emory Winship Cancer Institute (T.N.B.). We thank Emilee Wehunt and Haris Rashid, Clinical Research Coordinators in Emory’s Department of Neurosurgery, for help identifying GBM surgical cases and for obtaining pathology results for tissue samples. We thank Dr. Melissa Cadena, Dr. Maureen Sampson, and Dr. Emily Hill for helpful discussions on bioinformatic analyses and Dr. Youssef Zohdy for sharing some GBO lines used in this study. We also thank Dr. Jimena Andersen and the Andersen lab for helpful feedback on the use of spinal organoids and the rabies tracing technique. This research project was supported by the Emory University School of Medicine Flow Cytometry Core and the Winship Cancer Institute Cancer Center Support Grant P30CA138292.

## CONTRIBUTIONS

TNB and SAS designed all experiments and wrote the manuscript with input from all contributing authors. KH, EN, and JO provided all fresh surgical GBM sample resections used in this study. RDR created the gliomasphere cell lines used in this study. TNB performed all experiments, including organoid formation, GBM and organoid dissociations, engraftment experiments, immunohistochemistry, and transcriptomic analyses. TNB, SG, and MM performed infiltration assays and quantification. TNB and CS optimized the GBM procurement and dissociation protocol, developed the CNV analysis pipeline, and performed the CellChat analyses. AM and AB mapped scRNA-seq data onto the GBM meta-atlas. TNB, MM, and LN created and maintained GBO lines. TNB and URB performed the rabies tracing experiments. AS performed single-cell captures. AK and the BOH formed, maintained, and validated organoids.

---

**Fig. S1.**
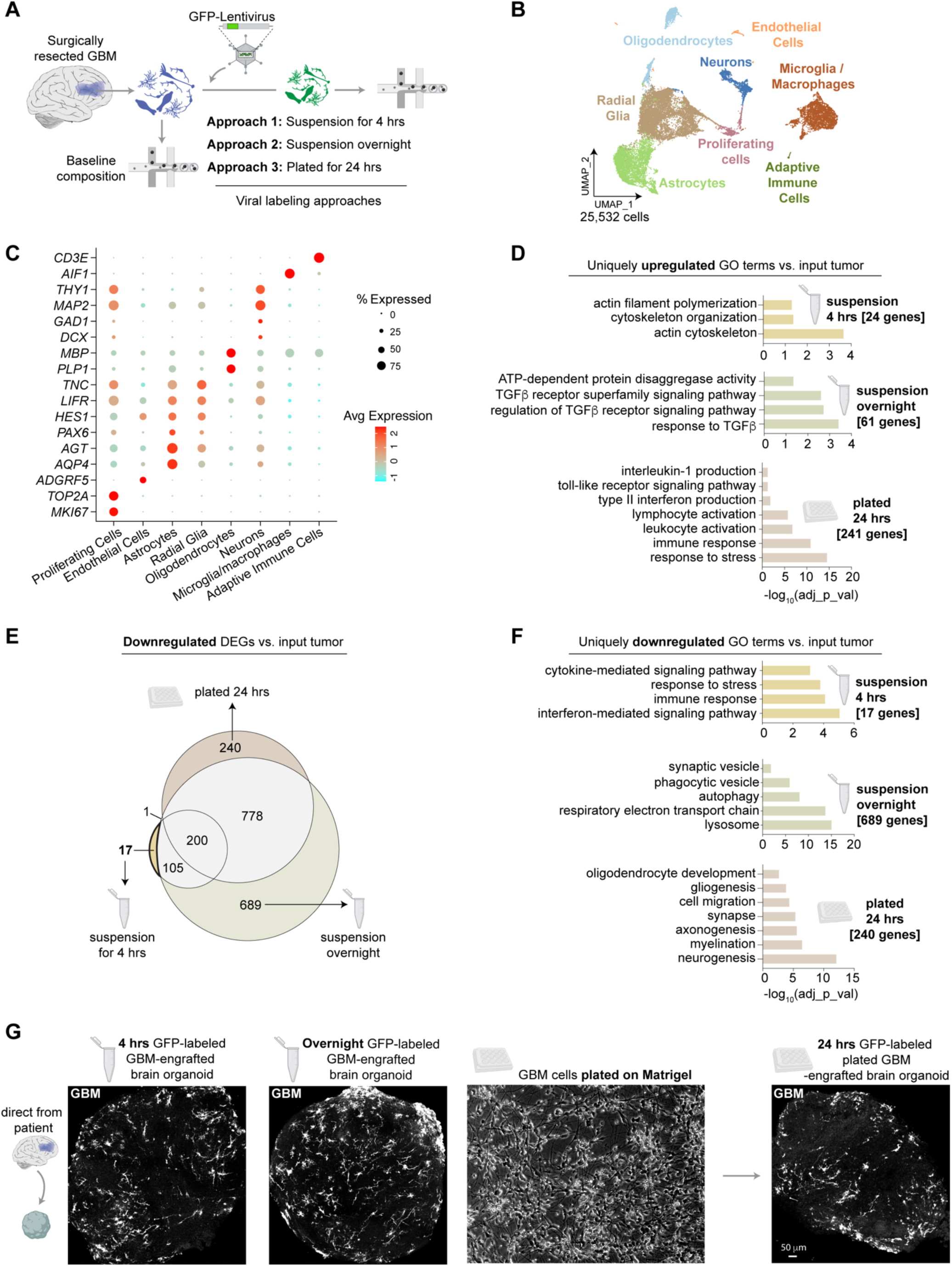
GFP-labeled GBM cells from surgically resected tumors infiltrate brain organoids regardless of lentiviral exposure or culture conditions. (**A**) Surgically resected GBM tissues were dissociated into a single-cell suspension and labelled with a GFP-lentivirus (pLenti-CMV-GFP) at a multiplicity of infection (MOI) of 5 by exposing cells to the virus for 4 hours in suspension culture on a shaker (Approach 1), overnight in suspension culture on a shaker (Approach 2), or by plating cells in 2D on Matrigel and then exposing cells to the virus for 24 hours (Approach 3). Single-cell RNA sequencing was performed following lentiviral exposure and compared to single-cell transcriptomes following dissociation of the input tumor. (**B**) UMAP of 25,532 individual cells that passed QC. (**C**) Expression of cell type-specific markers used to annotate the UMAP in b. (**D**) GO terms of genes uniquely upregulated across each of the 3 lentiviral exposure and culture conditions vs. the dissociated parent tumor. (**E**) Area-proportional Venn diagram to visualize the overlap between differentially downregulated genes when each of the lentiviral-exposed samples were compared to the dissociated, input tumor. The short, 4-hour GFP-lentivirus exposed GBM cells maintained in suspension culture showed the fewest differentially downregulated genes (17). (**F**) GO terms of genes uniquely downregulated across each of the 3 lentiviral exposure and culture conditions vs. the dissociated parent tumor. (**G**) Representative images of GBM (white)-engrafted brain organoid sections following 4 hours of GFP labeling in suspension culture, overnight GFP labeling in suspension culture, or after 24 hours of GFP labeling on GBM cells plated on Matrigel.

**Fig. S2.**
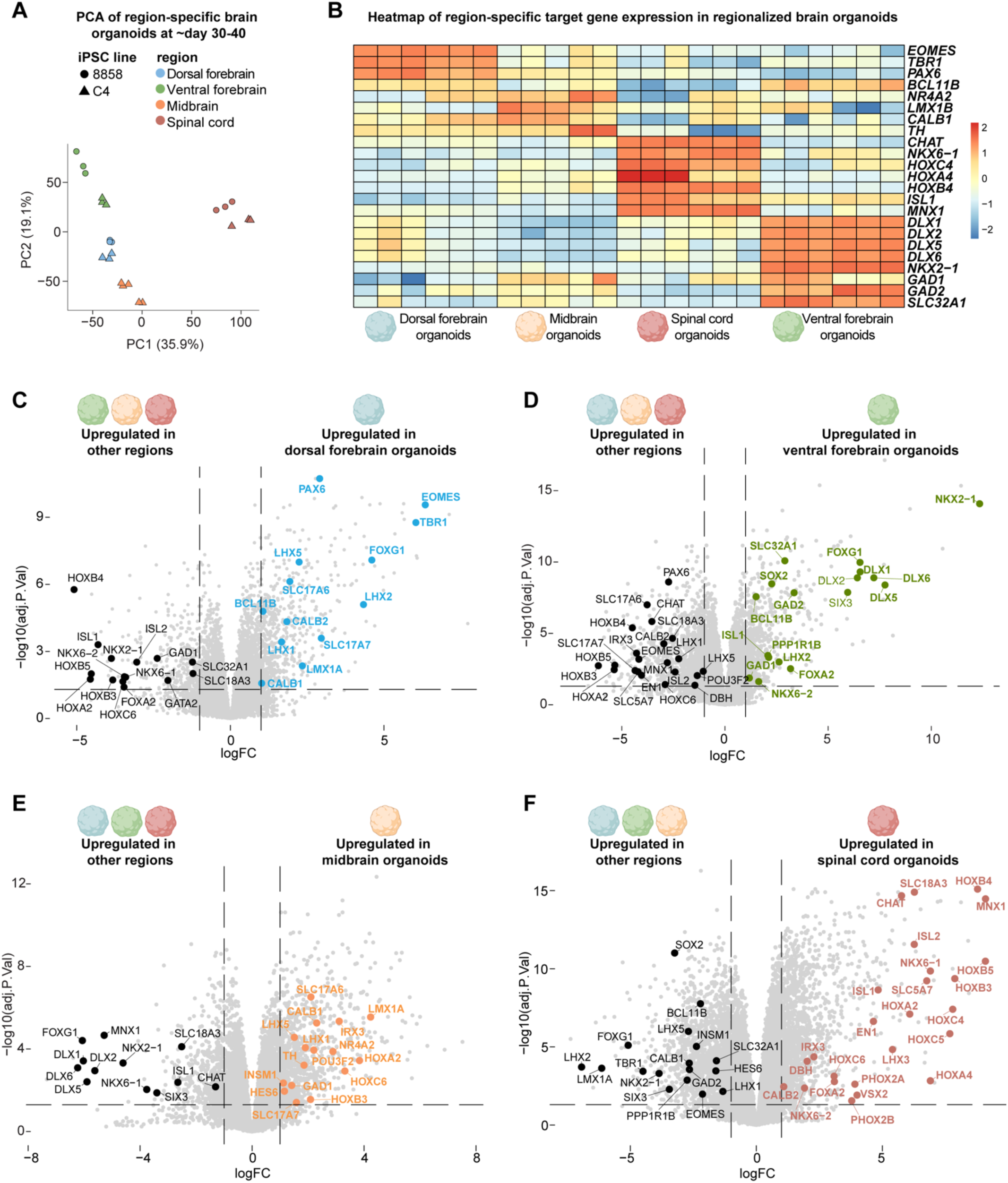
Validation of region-specific hiPSC-derived brain organoids at the transcriptomic level. (**A**) Principal component analysis (PCA) of bulk transcriptomes of hiPSC-derived dorsal forebrain, ventral forebrain, midbrain, and spinal cord organoids at ∼day 30-40. (**B**) Heatmap showing the expression of select region-specific genes across all four regionalized brain organoids. (**C**) Volcano plot showing differentially upregulated genes in dorsal forebrain organoids (right) vs. the other 3 regionalized organoids (left). (**D**) Volcano plot showing differentially upregulated genes in ventral forebrain organoids (right) vs. the other 3 regionalized organoids (left). (**E**) Volcano plot showing differentially upregulated genes in midbrain organoids (right) vs. the other 3 regionalized organoids (left). (**F**) Volcano plot showing differentially upregulated genes in spinal cord organoids (right) vs. the other 3 regionalized organoids (left).

**Fig. S3.**
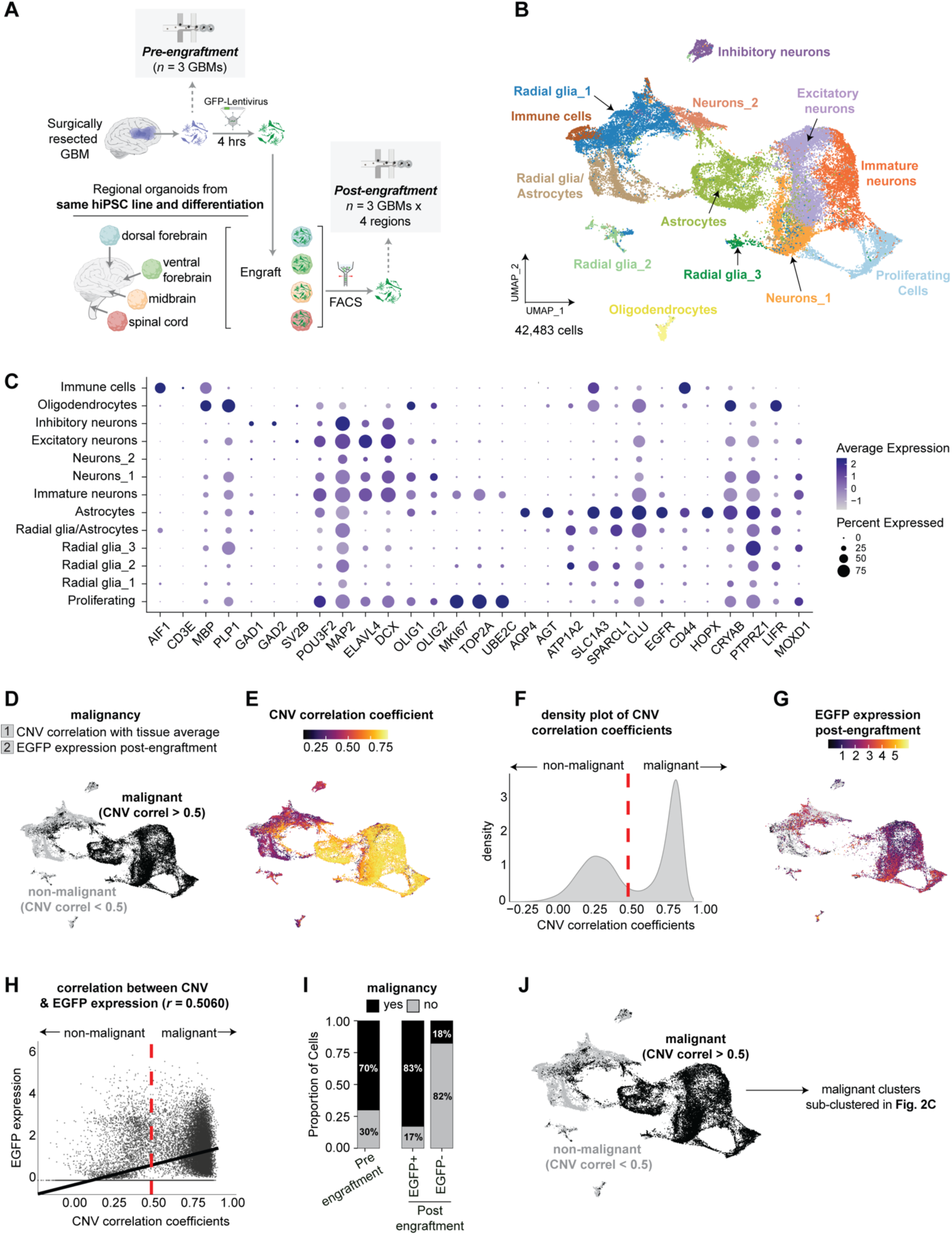
Identification of malignant clusters from single-cell RNA sequencing of direct-from-patient GBMs before and after engraftment into region-specific brain organoids. (**A**) GBM cells from surgical resections (*n=*3 independent patient samples) were dissociated, labeled with a GFP-lentivirus for 4 hours in suspension culture on a shaker, and then engrafted into dorsal forebrain, ventral forebrain, midbrain, and spinal cord organoids all derived in parallel from the same hiPSC lines and differentiations to reduce batch variability. After 14 days of organoid engraftment, GFP^+^ GBM cells were isolated via FACS and captured for single-cell RNA sequencing. For every post-engraftment GBM there was a matched pre-engraftment GBM to track the changes in the transcriptomic signatures of GBM cells after entry into the organoid niche. (**B**) UMAP of 42,483 individual cells that passed QC. (**C**) Expression of cell type-specific markers used to annotate the UMAP in b. (**D**) The inferCNV^92^ package was used to obtain copy number variation (CNV) signal for each cell, which was then correlated with the average CNV signal across all cells from the same tissue, as in our prior work^50^. Based on this method, we used a correlation coefficient of 0.5 as our cutoff to identify malignant cells. (**E**) Distribution of CNV correlation coefficients in the UMAP space. (**F**) Histogram of the density of cells below and above the CNV correlation coefficient cutoff of 0.5 reveals a bimodal distribution of cells. (**G**) The CNV correlation pipeline was validated by visualizing the expression of *EGFP* post-engraftment in the UMAP space. (**H**) The expression of *EGFP* post-engraftment was correlated with the CNV coefficients. (**I**) Proportion of cells annotated as malignant vs. non-malignant before and after engraftment. (**J**) Distribution of malignant cells (CNV correlation coefficient > 0.5) and non-malignant cells (CNV correlation coefficient < 0.5) visualized in the UMAP space. The malignant cells were then subclustered and shown in Fig. 2C.

**Fig. S4.**
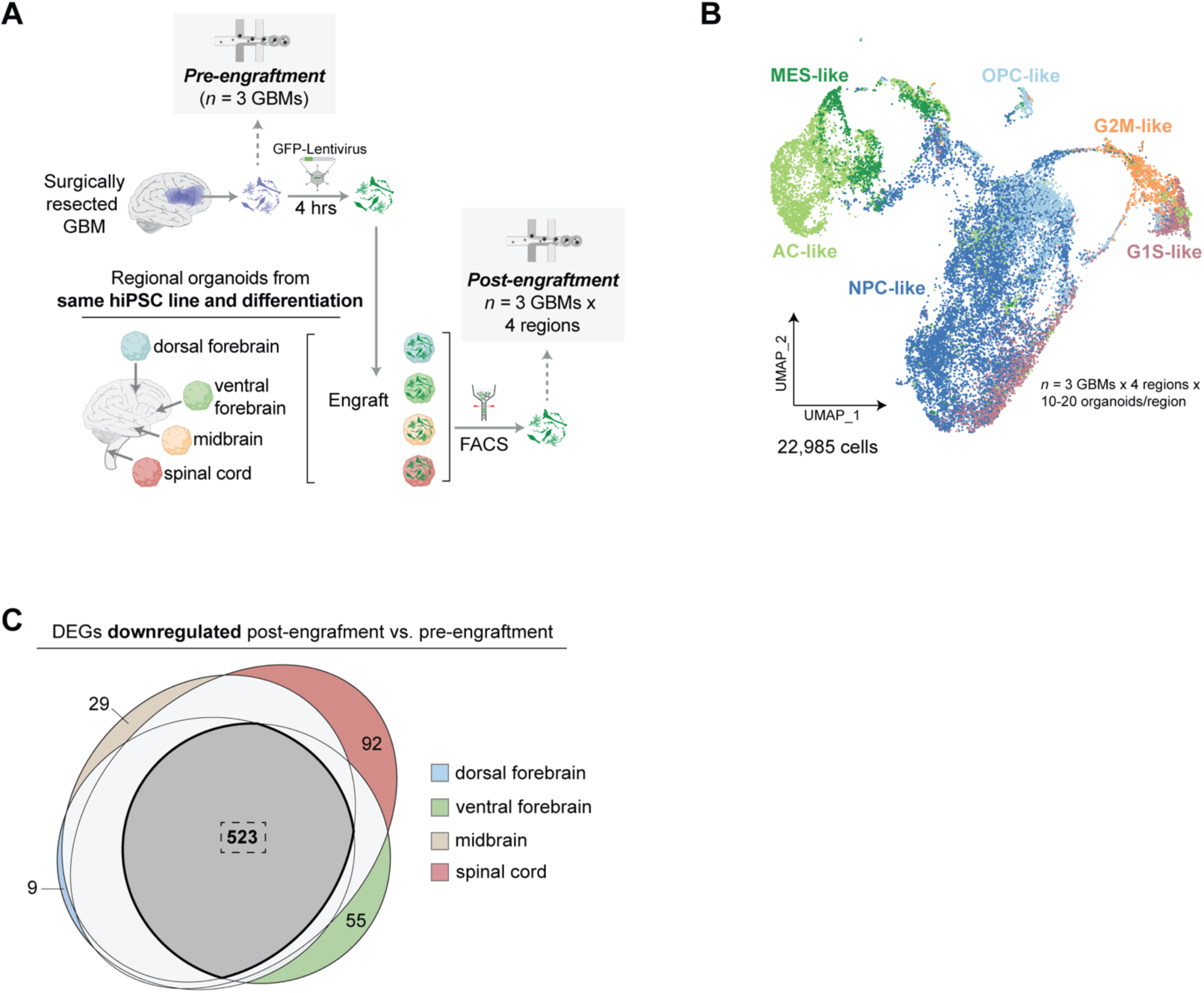
GBM cells downregulate similar transcriptional signatures post-engraftment into region-specific brain organoids. (**A**) GBM cells from surgical resections (*n=*3 independent patient samples) were dissociated, labeled with a GFP-lentivirus for 4 hours in suspension culture, and then engrafted into dorsal forebrain, ventral forebrain, midbrain, and spinal cord organoids all derived in parallel from the same hiPSC lines and differentiations to reduce batch variability. After 14 days of organoid engraftment, GFP^+^ GBM cells were isolated via FACS and captured for single-cell RNA sequencing. For every post-engraftment GBM there was a matched pre-engraftment GBM to track the changes in the transcriptomic signatures of GBM cells after entry into the organoid niche. (**B**) UMAP of 22,985 individual malignant cells that passed QC and after exclusion of non-malignant cells based on copy number variations. Malignant cells were assigned to clusters based on Neftel *et. al.*, 2019^21^. (**C**) Area-proportional Venn diagram to visualize the overlap between differentially downregulated genes when GBM cells engrafted into each regional organoid were compared to the dissociated tumor pre-engraftment. A majority of differentially downregulated genes (523) were shared across all four regional microenvironments. Select GO terms from gProfiler reflecting the 523 downregulated genes are shown in Fig. 2D.

**Fig. S5.**
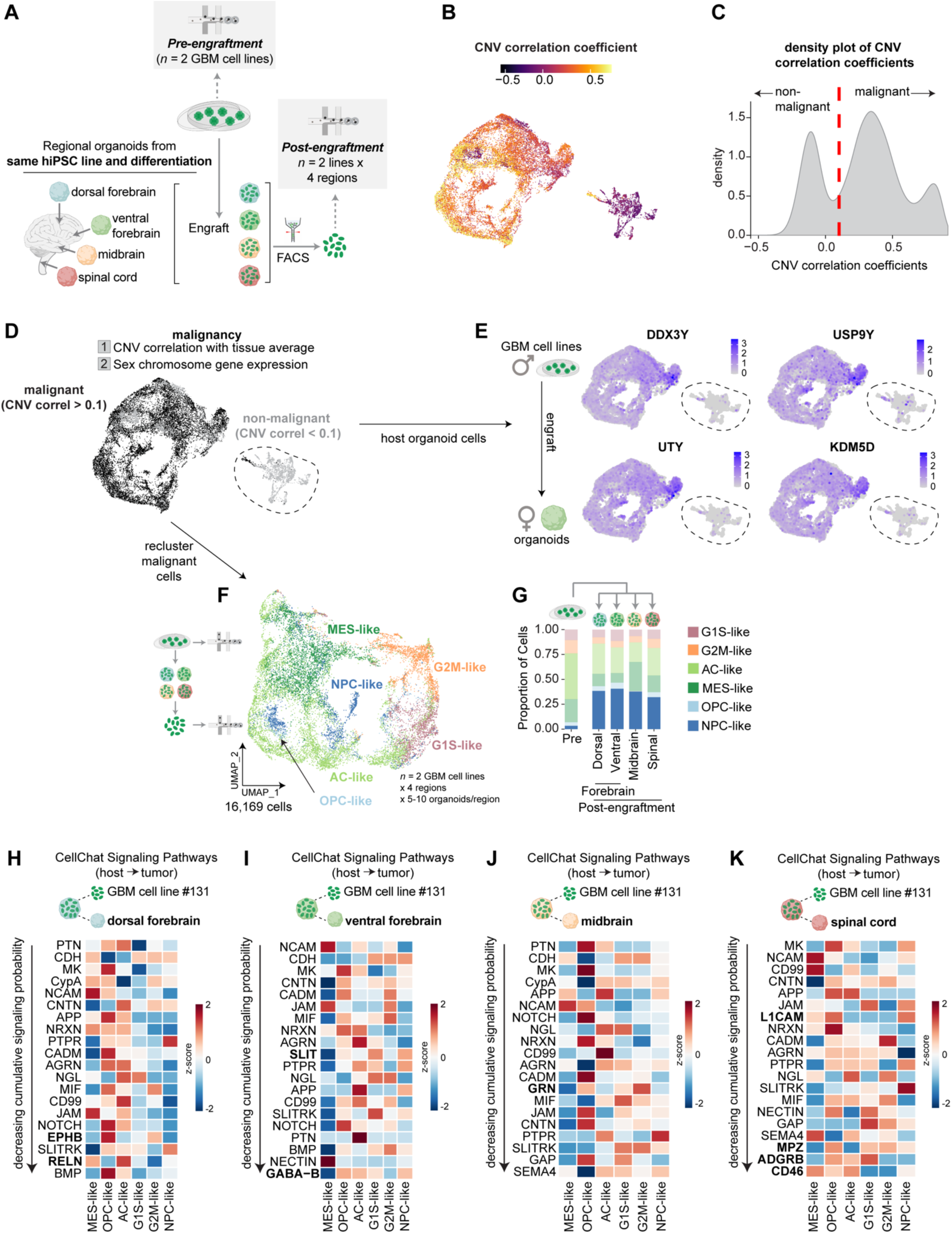
GBM cell lines are enriched in NPC-like transcriptomic signatures post-engraftment and may rely on shared extrinsic cues across brain regions. (**A**) GBM cells from GFP+ patient-derived cell lines (*n=*2 independent cell lines) were dissociated and engrafted into dorsal forebrain, ventral forebrain, midbrain, and spinal cord organoids all derived in parallel from the same hiPSC lines and differentiations to reduce batch variability. After 14 days of organoid engraftment, GFP^+^ GBM cells were isolated via FACS and captured for single-cell RNA sequencing. For every post-engraftment GBM there was a matched pre-engraftment GBM cell line to track the changes in the transcriptomic signatures of GBM cells after entry into the organoid niche. (**B**) Distribution of copy number variation (CNV) correlation coefficients, calculated as the correlation between the CNV signal of each cell and the average CNV signal across all cells from the same tissue, visualized in the UMAP space. (**C**) Histogram of the density of cells below and above the CNV correlation coefficient cutoff of 0.1 reveals a bimodal distribution of cells. (**D**) Distribution of malignant cells (CNV correlation coefficient > 0.1) and non-malignant cells (CNV correlation coefficient < 0.1) visualized in the UMAP space. (**E**) As male GBM cell lines were engrafted into organoids derived from a female hiPSC line, we could also visualize that the cells annotated as non-malignant (CNV correlation coefficient < 0.1) were host organoid cells, as they did not express the Y-chromosome genes *DDX3Y*, *USP9Y*, *UTY*, and *KDM5D*. (**F**) The malignant cells (CNV correlation coefficient > 0.1) were re-clustered and then annotated based on the method in Neftel *et. al.,* 2019^21^. (**G**) Proportion of cells across the six Neftel GBM cell types in in each of the four region-specific brain organoids and in the parent tumor. (**H**) CellChat analyses^57^ to identify ligand-receptor relationships between engrafted GBM cells and host organoid cells in the dorsal forebrain, arranged in decreasing order of the cumulative strength of their interaction. (**I**) CellChat analyses to identify ligand-receptor relationships between engrafted GBM cells and host organoid cells in the ventral forebrain, arranged in decreasing order of the cumulative strength of their interaction. (**J**) CellChat analyses to identify ligand-receptor relationships between engrafted GBM cells and host organoid cells in the midbrain, arranged in decreasing order of the cumulative strength of their interaction. (**K**) CellChat analyses to identify ligand-receptor relationships between engrafted GBM cells and host organoid cells in the spinal cord, arranged in decreasing order of the cumulative strength of their interaction. Ligands in **H-K** that are unique to each region are written in bold.

**Fig. S6.**
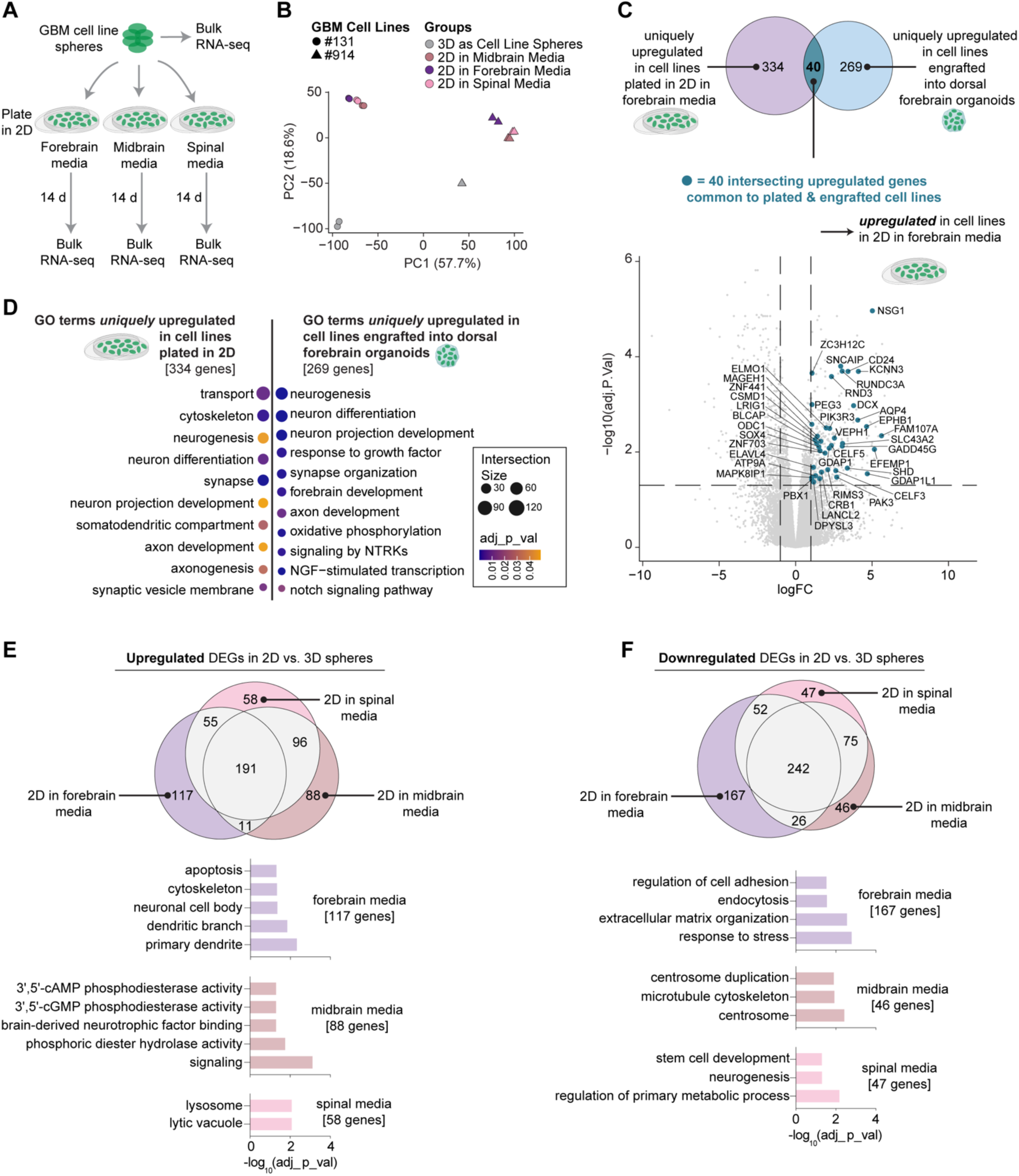
Transcriptomic changes following engraftment of GBM cell lines into brain organoids are largely distinct from those observed in 2D adherent culture. **(A)** GBM cell line spheres (*n=*2 independent cell lines) were subject to bulk RNA sequencing before and after 14 days of adherent culture in forebrain media (neurobasal), midbrain media (neurobasal, supplemented with BDNF, GDNF, cAMP, and L-ascorbic acid), and spinal cord media (neurobasal, supplemented with N2, BDNF, cAMP, L-ascorbic acid, and IGF). (**B**) Principal component analysis (PCA) reveals that the primary variation is a result of cell line (PC1 = 57.7%), followed by culture conditions (3D spheres vs. 2D adherent culture; PC2 = 18.6%). (**C**) Area-proportional Venn diagram to visualize the overlap between differentially upregulated genes when GBM cell lines are plated in 2D (vs. 3D spheres; bulk transcriptomes) and when GBM cell lines are engrafted into dorsal forebrain organoids (vs. pre-engrafted 3D spheres; single-cell transcriptomes). Only a small proportion (40) of genes were overlapping, revealing largely unique gene expression patterns. Volcano plot (bottom of **C**) shows the 40 genes that are commonly upregulated, irrespective of whether GBM cell lines are engrafted into organoids or plated in 2D. (**D**) GO terms related to genes that are uniquely upregulated when GBM cell lines are plated in 2D (left; 334 unique genes) vs. when GBM cell lines are engrafted into dorsal forebrain organoids (right; 269 unique genes). (**E**) Area-proportional Venn diagram to visualize the overlap between differentially upregulated genes when GBM cell lines are plated in 2D in forebrain media, midbrain media, and spinal cord media. GO terms related to genes that are uniquely upregulated in each media condition are shown below. (**F**) Area-proportional Venn diagram to visualize the overlap between differentially downregulated genes when GBM cell lines are plated in 2D in forebrain media, midbrain media, and spinal cord media. GO terms related to genes that are uniquely downregulated in each media condition are shown below.

**Fig. S7.**
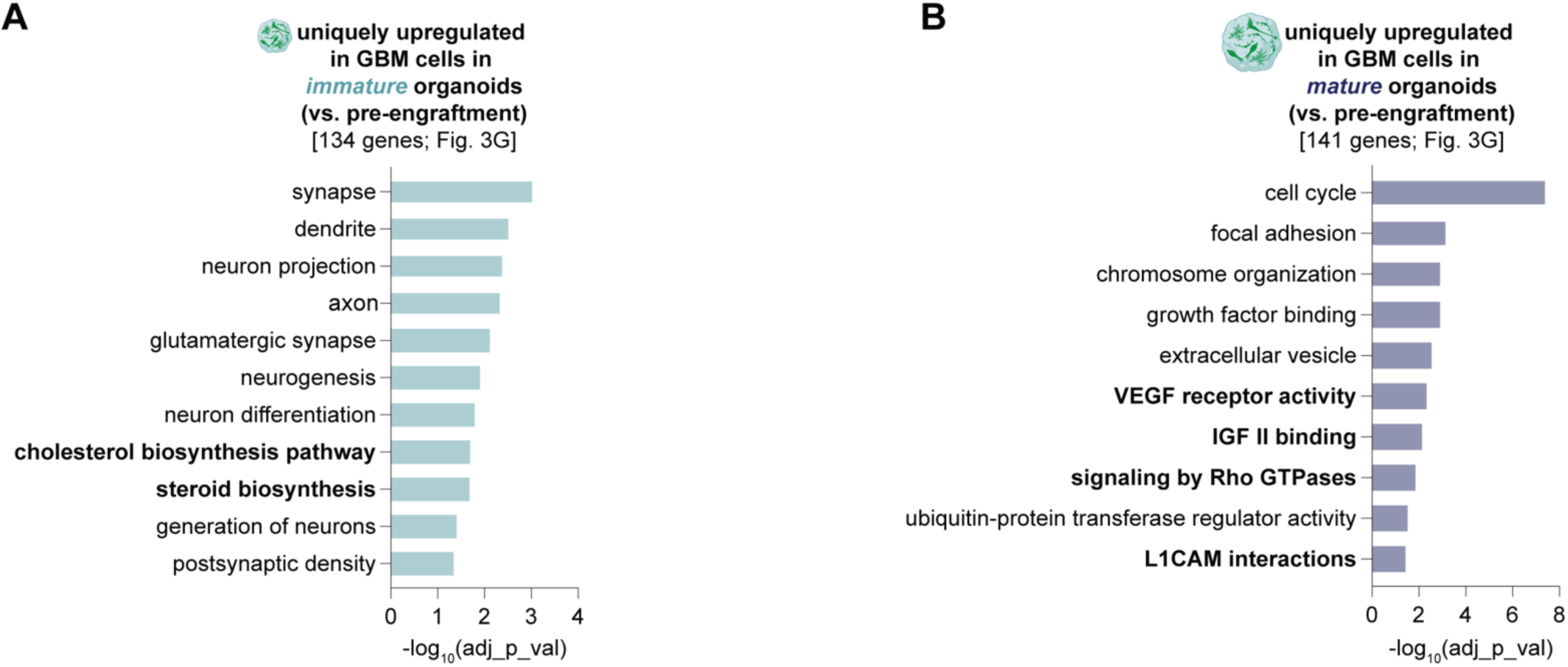
GO terms reflecting genes uniquely upregulated when direct-from-patient GBM cells are engrafted into immature organoids and mature organoids. (**A**) GO terms that are over-represented when examining the 134 genes that are uniquely upregulated when GBM cells from surgical resections are engrafted into immature dorsal forebrain organoids (from the Venn diagram in Fig. 3G). (**B**) GO terms that are over-represented when examining the 141 genes that are uniquely upregulated when GBM cells from surgical resections are engrafted into mature dorsal forebrain organoids (from the Venn diagram in Fig. 3G).

**Fig. S8.**
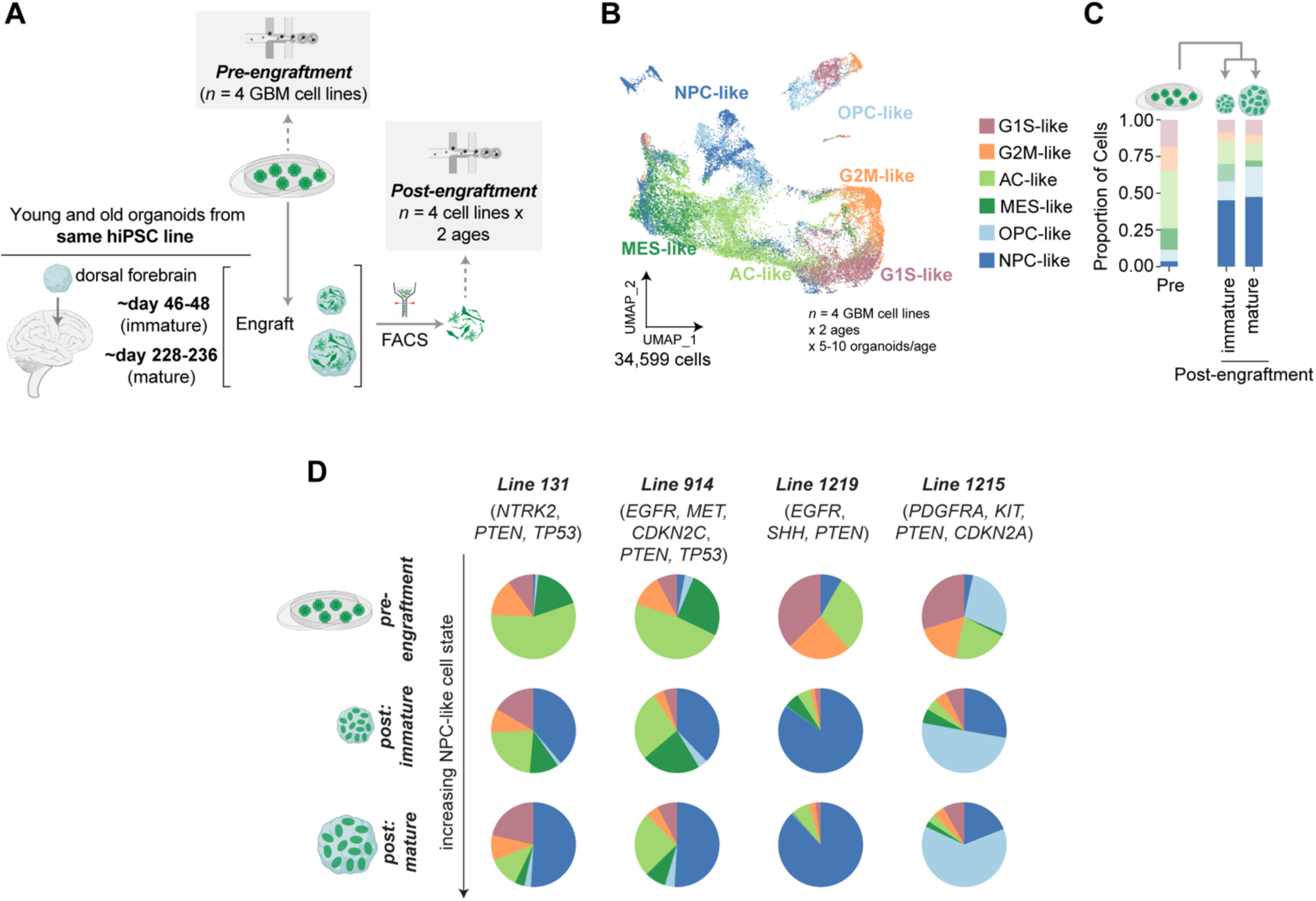
GBM cell lines are also enriched in NPC-like transcriptomic signatures post-engraftment into mature (day 200+) dorsal forebrain organoids. (**A**) GBM cells from GFP+ patient-derived cell lines (*n=*4 independent cell lines) were dissociated and engrafted into dorsal forebrain organoids that were immature (∼day 46-48 on the day of engraftment) vs. dorsal forebrain organoids that were more mature (∼day 228-236). After 14 days of organoid engraftment, GFP^+^ GBM cells were isolated via FACS and captured for single-cell RNA sequencing. For every post-engraftment GBM there was a matched pre-engraftment GBM cell line to track the changes in the transcriptomic signatures of GBM cells after entry into the organoid niche. (**B**) UMAP of 34,599 individual malignant cells that passed QC and annotated as in Neftel *et. al.,* 2019. (**C**) Proportion of cells across the six Neftel GBM cell types in immature and more mature dorsal forebrain organoids, and in the parent GBM cell line. (**D**) Distribution of cells across each of the six Neftel GBM cell types, visualized as pie charts and separated based on the genetic background of the GBM cell line.

**Fig. S9.**
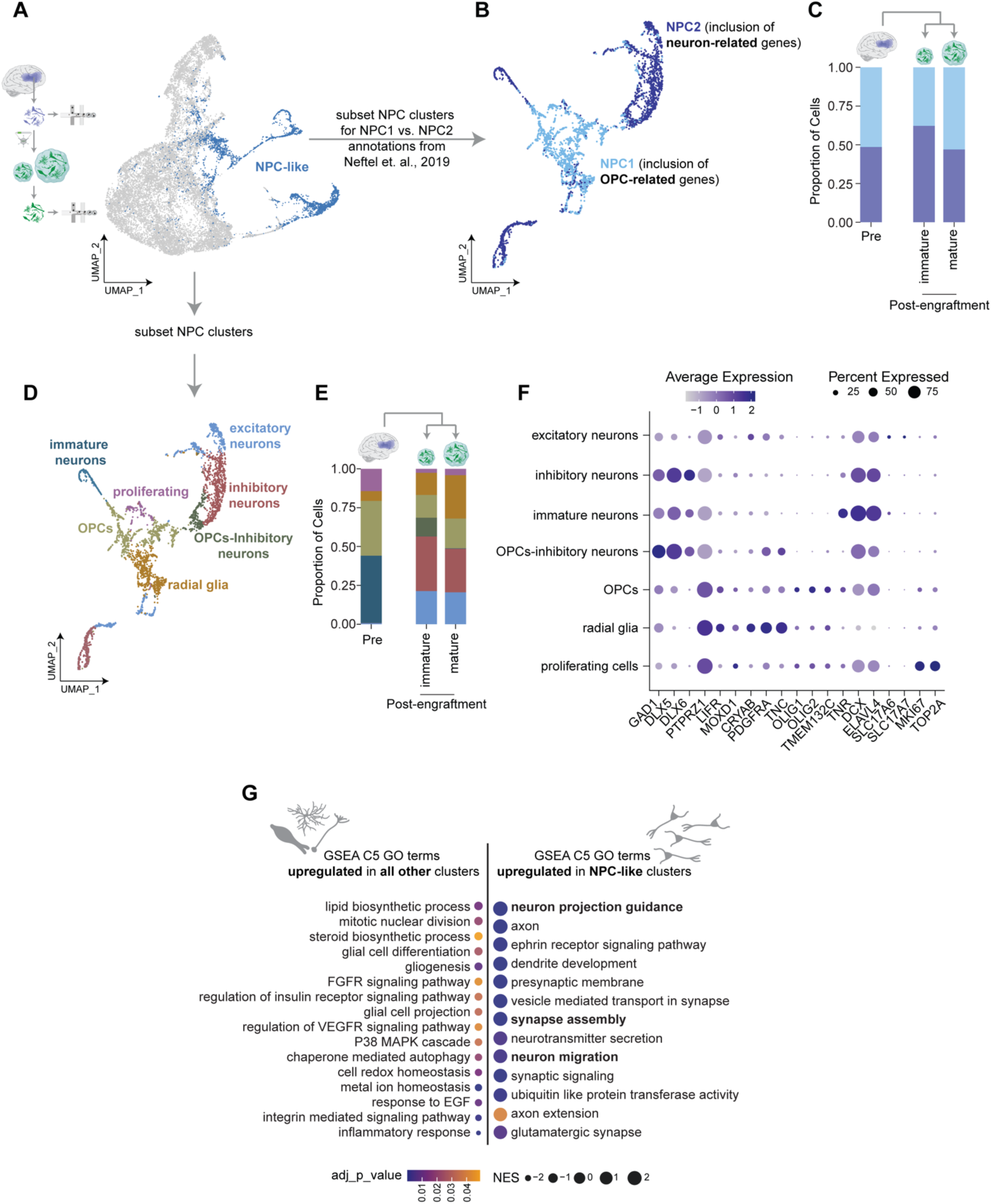
The NPC-like transcriptomic state largely reflects neuronal gene expression signatures. (A) NPC-like cells from Fig. 3 were subclustered to further resolve the heterogeneity of this GBM cell type. (B) NPC-like GBM cells annotated by whether they co-express neuron-like genes (NPC2 subtype from Neftel *et. al.*) or oligodendrocyte precursor (OPC)-like genes (NPC1 subtye from Neftel *et. al.*). (**C**) Proportion of NPC1 and NPC2 GBM cells pre-engraftment and after engraftment into immature and more mature dorsal forebrain organoids. (**D**) NPC-like GBM cells manually annotated by cell type-specific gene expression. (**E**) Proportion of NPC-like GBM cells that reflect proliferating signatures, OPC-like signatures, radial glia-like signatures, immature neuron-like signatures, excitatory neuron-like signatures, and inhibitory neuron-like signatures, before and after engraftment into immature and more mature dorsal forebrain organoids. (**F**) Dot plot showing cell type-specific genes used to annotate the UMAP in d. (**G**) Gene set enrichment analyses (GSEA) analyses reflecting GO terms that are over-represented in NPC-like clusters (right) vs. all other Neftel clusters (left).

**Fig. S10.**
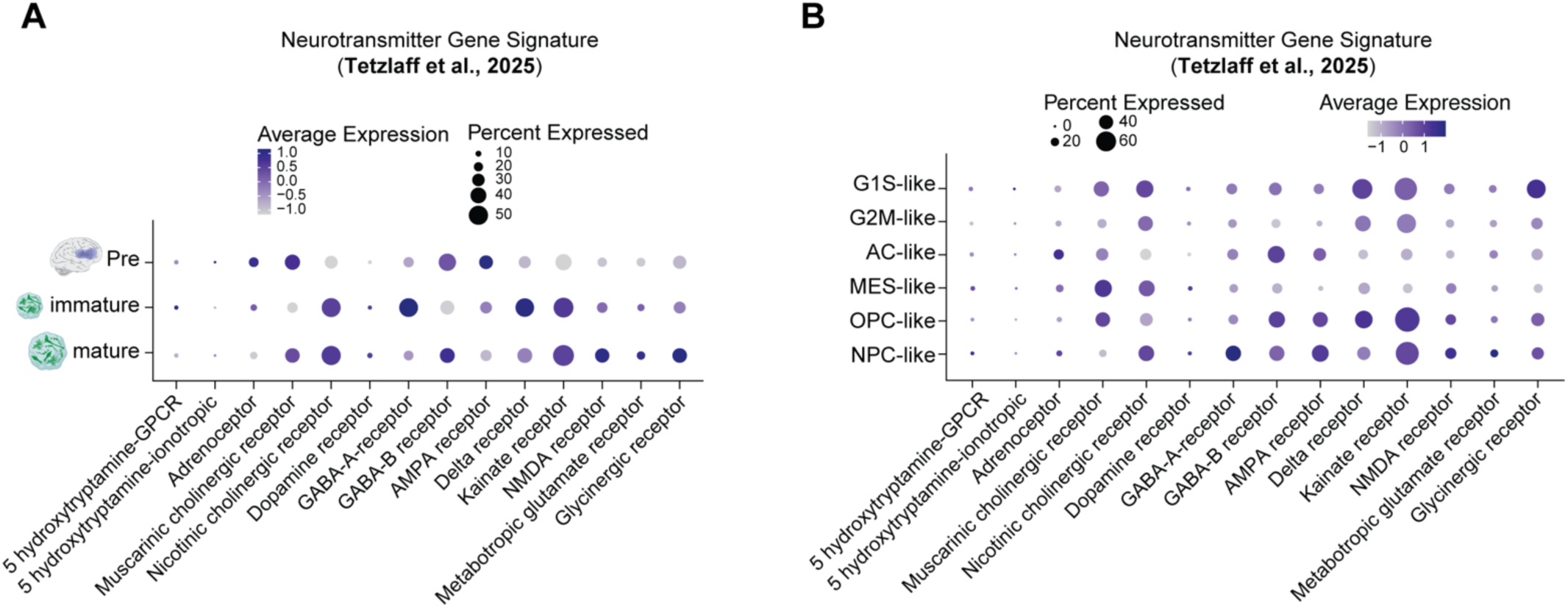
Expression of neurotransmitter receptor genes in GBM cells before and after engraftment into immature and more mature dorsal forebrain organoids. (**A**) Neurotransmitter receptor gene expression signatures^54^, displayed for GBM cells before vs. after organoid engraftment into immature and more mature dorsal forebrain organoids, shown here using the transcriptomic data from Fig. 3. (**B**) Neurotransmitter receptor gene expression signatures^54^, shown here based on the Neftel *et. al.* cluster annotations of the transcriptomic data from Fig. 3.

**Fig. S11.**
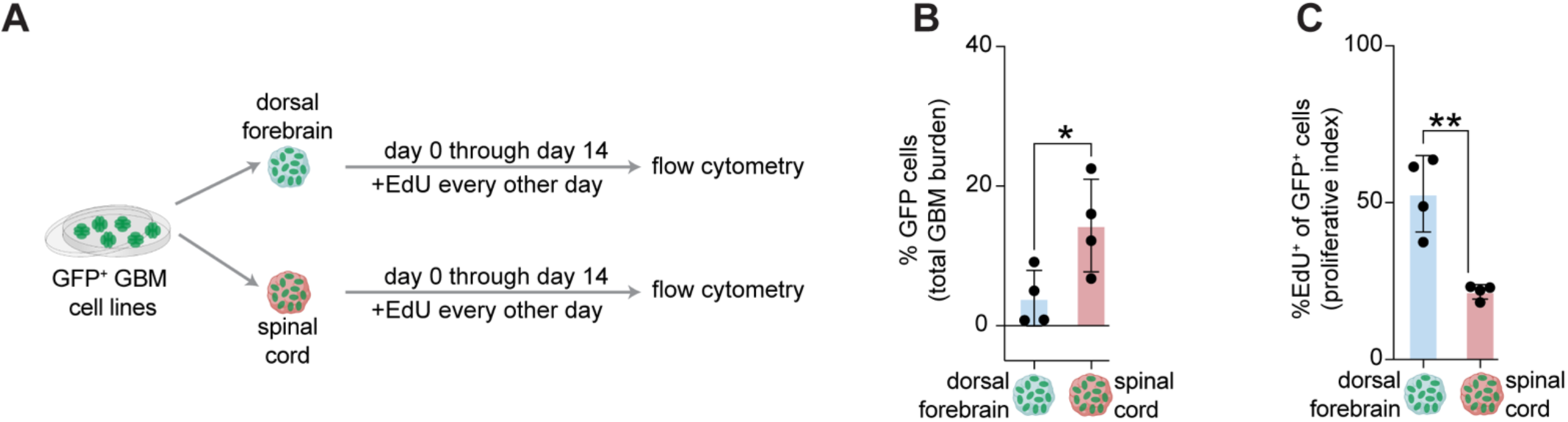
GBM cell lines are more proliferative in dorsal forebrain organoids than spinal cord organoids. (**A**) GFP+ GBM cells from patient-derived cell lines were engrafted into dorsal forebrain and spinal cord organoids in parallel and then exposed to the thymidine analogue EdU during the 14-day engraftment period. (**B**) Assessment of total burden across each organoid niche, measured via FACS. (**C**) Proliferation of GFP^+^ GBM cells in each organoid niche, measured via FACS. In b-c, data are from two independent GBM cell lines engrafted into two independent hiPSC-derived brain organoid lines (*n=*2 GBM lines x 2 organoid lines). Data were normally distributed (based on Shapiro-Wilk) and analyzed by the two-tailed unpaired *t*-test. All error bars are mean + SD.

**Fig. S12.**
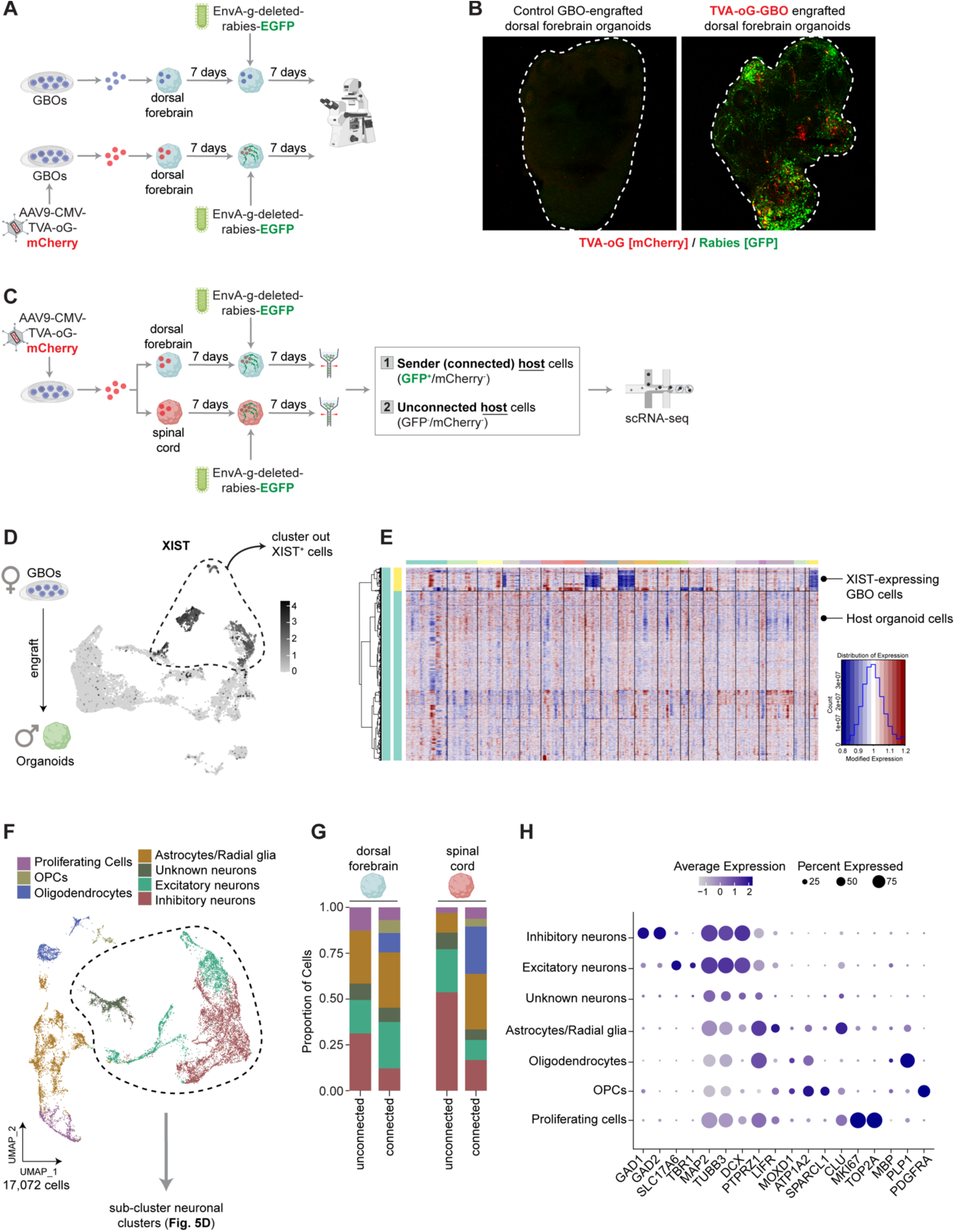
Host organoid cells synaptically connected to GBM are less likely to be inhibitory neurons. (**A**) Whole GBM organoids (GBOs) were either maintained as controls or transduced with AAV9-CMV-TVA-mCherry-2A-oG, following which GBM cells were engrafted into dorsal forebrain and spinal cord organoids. After 7 days, engrafted organoids were infected with EnvA G-deleted rabies-EGFP and 7 days later (14 days post-engraftment) organoids were fixed for immunohistochemistry. (**B**) Representative image shows that the rabies virus (green) selectively infects cells in the TVA-oG-GBO engrafted brain organoids, but not in control GBO engrafted brain organoids. (**C**) Synaptically connected host organoid cells (GFP^+^ / mCherry^-^) and unconnected host organoid cells (GFP^-^ / mCherry^-^) in the dorsal forebrain and spinal cord were isolated by FACS and subject to single-cell captures. (**D**) As a female GBO line was engrafted into brain organoids derived from a male hiPSC line, we used *XIST* to selectively cluster out cells that were likely GBM cells. (**E**) Heatmap of copy number variation (CNV) signal confirms that *XIST*-expressing cells are malignant GBM cells. (**F**) Non-malignant host organoid cells were manually annotated using gene expression of cell type-specific markers and revealed the presence of proliferating cells, OPCs, oligodendrocytes, astrocytes/radial glia, and neurons. (**G**) Proportion of cells across each of the annotated cell types in the synaptically connected and unconnected host fractions in the dorsal forebrain and spinal cord. (**H**) Dot plot showing cell type-specific genes used to annotate the UMAP in f. The neuronal clusters in f were re-clustered (and shown in Fig. 5D) to enhance granularity and identify distinct subclusters of excitatory and inhibitory neurons.

**Fig. S13.**
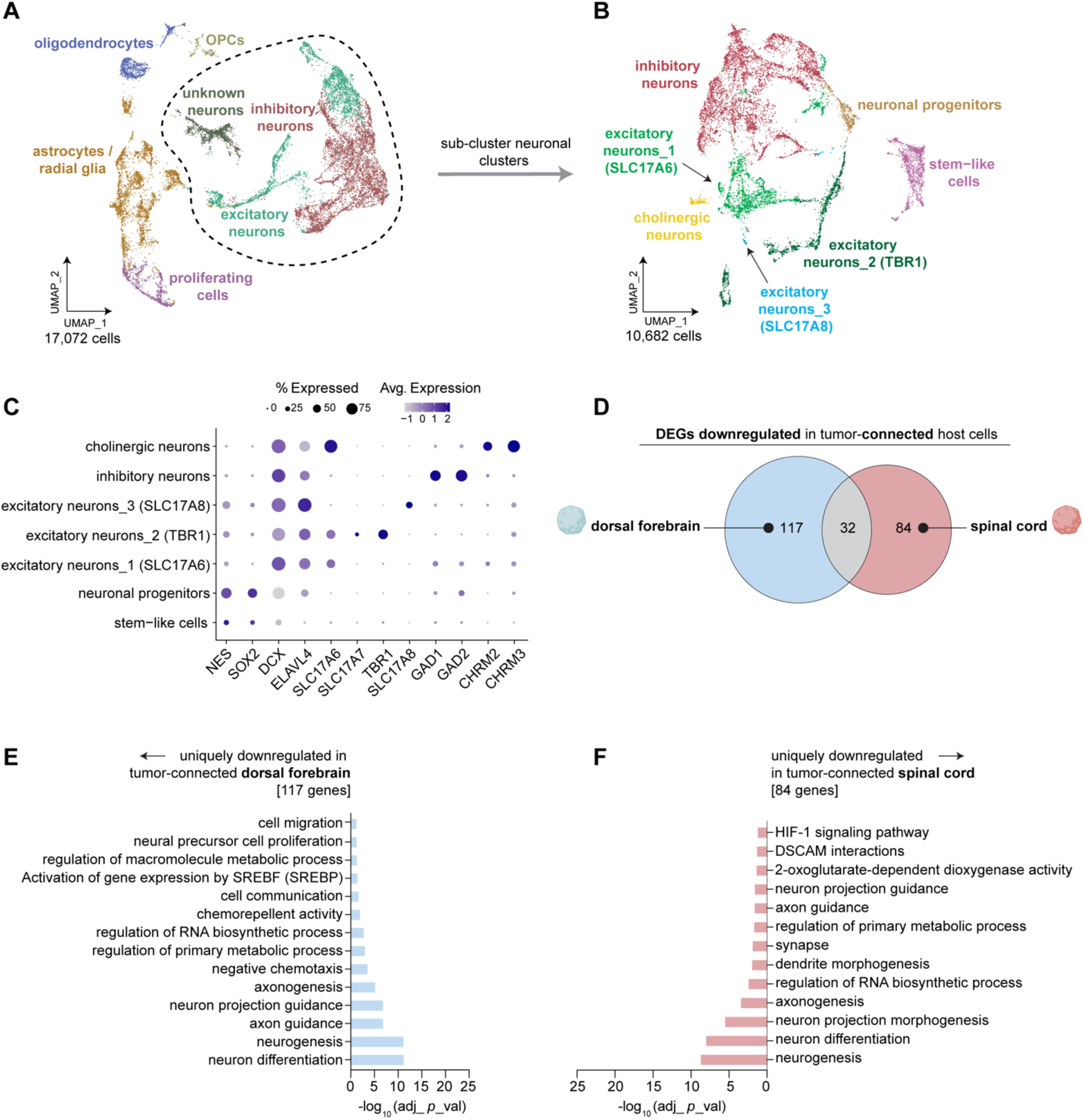
Genes that are downregulated in host cells synaptically connected to GBM are largely distinct in the dorsal forebrain and the spinal cord. (**A**) Neuron-related clusters from single-cell captures following rabies-based tracing in GBM-engrafted dorsal forebrain vs. spinal cord organoids (from Fig. 5 and Fig. S12) were re-clustered. (**B-C**) UMAP manually annotated with identities of neurons, based on expression of neuronal subtype-specific markers. (**D**) Area-proportional Venn diagram to visualize the overlap between differentially downregulated genes when host cells that are synaptically connected to GBM are compared to the host cells synaptically unconnected to GBM in the dorsal forebrain (blue) vs. the spinal cord (red) microenvironments. (**E**) GO terms related to the 117 genes that are uniquely downregulated in the GBM-connected host fraction in the dorsal forebrain. (**F**) GO terms related to the 84 genes that are uniquely downregulated in the GBM-connected host fraction in the spinal cord.

